# Spastin locally amplifies microtubule dynamics to pattern the axon for presynaptic cargo delivery

**DOI:** 10.1101/2023.08.08.552320

**Authors:** Jayne Aiken, Erika L. F. Holzbaur

## Abstract

Neurons rely on long-range trafficking of synaptic components to form and maintain the complex neural networks that encode the human experience. With a single neuron capable of forming thousands of distinct *en passant* synapses along its axon, spatially precise delivery of the necessary synaptic components is paramount. How these synapses are patterned, and how efficient delivery of synaptic components is regulated, remains largely unknown. Here, we reveal a novel role for the microtubule severing enzyme spastin in locally enhancing microtubule polymerization to influence presynaptic cargo pausing and retention along the axon. In human neurons derived from induced pluripotent stem cells (iPSCs), we identify sites stably enriched for presynaptic components, termed ‘protosynapses’, which are distributed along the axon prior to the robust assembly of mature presynapses apposed by postsynaptic contacts. These sites are capable of cycling synaptic vesicles, are enriched with spastin, and are hotspots for new microtubule growth and synaptic vesicle precursor (SVP) pausing/retention. Disruption of neuronal spastin, either by CRISPRi-mediated depletion or transient overexpression, interrupts the localized enrichment of dynamic microtubule plus ends and diminishes SVP accumulation. Using an innovative human heterologous synapse model, where microfluidically isolated human axons recognize and form presynaptic connections with neuroligin-expressing non-neuronal cells, we reveal that neurons deficient for spastin do not achieve the same level of presynaptic component accumulation as control neurons. We propose a model where spastin acts locally as an amplifier of microtubule polymerization to pattern specific regions of the axon for synaptogenesis and guide synaptic cargo delivery.

## INTRODUCTION

The human central nervous system is comprised of billions of neurons that connect with one another through trillions of synapses. With a single neuron responsible for actively maintaining potentially thousands of individual pre- and postsynaptic connections, a considerable challenge is faced during neurodevelopment: where do individual synapses form and how are they maintained?

Neurons rely on long-range, microtubule-based trafficking to supply components necessary for both initial synapse set-up and continued upkeep. As neurons contain distinct compartments including the soma, dendrites (hosting postsynapses), and the axon (hosting presynapses), the sorting, transport, and appropriate delivery of cargo must be stringently regulated. Microtubules are polarized cytoskeletal polymers that provide structural integrity to axons and dendrites and serve as the ‘tracks’ upon which motor proteins transport cargo. In the axon, microtubules are organized with their dynamic plus ends facing out toward the axon tip [1]. Anterograde axonal movement (from the soma out toward the axon tip) is completed by kinesin motors, which move processively toward microtubule plus ends. Retrograde movement (from the distal axon back to the soma) is performed by dynein motors, which move toward microtubule minus ends. These key molecular players of long-distance transport must undergo complex, circumstance-specific regulation to sustain neuronal health and connectivity [2, 3].

The cellular mechanisms dictating how and where synapses are initiated are not well understood. Classical work suggests a temporal progression from initial recognition of synaptic partners, assembly of pre- and postsynaptic compartments, and finally the maturation of fully functional synapses, all of which rely on the directed trafficking of synaptic components. However, there is evidence for both pre- and postsynaptic component accumulation and function prior to contact with a synaptic partner. This has been observed *in vivo* in *Drosophila* motor neurons, which form morphologically normal presynaptic active zones independent of postsynaptic target cells [4], as well as in cultured mammalian neurons, where presynaptic machinery capable of cycling synaptic vesicles is observed irrespective of postsynaptic contacts [5]. Further, dendritic filopodia preferentially form stable contacts with nascent presynaptic sites [6], suggesting that pre-patterning of synaptic components can influence synapse site selection.

Once presynapses have formed, recent work demonstrates that local microtubule dynamics influence synaptic vesicle precursor (SVP) pausing, retention, and activity-induced exchange between boutons [7, 8]. SVPs are synthesized in the soma, transported along the axon by the kinesin-3 motor KIF1A [9], and eventually captured at presynaptic sites along the axon, termed *en passant* synapses, where they mature into synaptic vesicles [10]. In mature primary rat hippocampal neurons, *en passant* synaptic regions are hotspots for new microtubule polymerization [7]. This local enrichment of microtubule growth directly influences SVP pausing and retention as KIF1A preferentially detaches from GTP-rich microtubule plus ends [7]. Strikingly, anterograde SVP retention at *en passant* synapses can be curtailed either by reducing microtubule polymerization or through a KIF1A mutation that diminishes microtubule plus-end detachment [7]. Together, these observations indicate a mechanistic link between local microtubule dynamics and KIF1A-mediated SVP delivery to presynaptic regions. Which factors locally regulate new microtubule growth at presynaptic zones along the axon has yet to be investigated.

Paradigm-shifting studies using reconstituted single-molecule assays have revealed that the microtubule severing enzyme spastin, traditionally thought to dismantle microtubule networks, can also act to amplify microtubule mass *in vitro* [11, 12]. Spastin is a hexameric AAA-ATPase enzyme that uses energy obtained from ATP hydrolysis to cut microtubules into shorter filaments by pulling tubulin heterodimers from the lattice [13–15]. Early studies in *Drosophila* exposed the paradoxical nature of spastin’s microtubule severing activity *in vivo*: spastin overexpression was found to dismantle microtubule networks in muscle cells, but instead of observing increased microtubule mass in *spastin*-null mutants, fewer microtubule bundles were found in the neuromuscular junction [13]. These apparently conflicting cellular data have been recapitulated in different model systems and cell types, revealing reduction of microtubule polymerization and/or mass upon spastin overexpression, knockdown, or mutation [13, 16–19]. In motor neurons, spastin enrichment has been observed in axonal growth cones [20], axonal branch sites [20], and neuromuscular synaptic boutons [13]. Spastin’s potential role in regulating synaptic connections of the central nervous system (CNS) is bolstered by the observation that spastin knockout mice exhibit fewer hippocampal synapses [21]. Further, human patients harboring spastin mutations frequently present with psychiatric comorbidities including memory impairment, intellectual disability, autism spectrum disorder, and severe depression [22]. Together, these observations raise the compelling possibility that spastin may regulate synapse biology by influencing microtubule dynamics.

Here, we find that spastin acts as a local amplifier of microtubule polymerization to pattern the axon for presynaptic cargo accumulation in human iPSC-derived neurons. Prior to the formation of robust synaptic connections, we note the accumulation of stable populations of presynaptic components along the axon. These sites are hotspots for dynamic microtubules and are marked by SVP pausing in both the anterograde and retrograde directions. Spastin depletion leads to decreased microtubule polymerization and SVP pauses/retention at presynaptic accumulation sites, while spastin overexpression leads to increased microtubule polymerization mislocalized outside of these zones. Thus, while spastin has been canonically considered to disassemble microtubule networks [13, 23], we find instead that spastin activity amplifies dynamic microtubule plus ends to locally regulate presynaptic trafficking. To further interrogate spastin’s role in presynapse formation, we developed a heterologous synapse assay in which microfluidically isolated human axons rapidly induce robust presynapse formation upon contact with neuroligin-expressing HEK cells. This assay reveals that axonal spastin enhances delivery of presynaptic components. These findings support a model where spastin locally enhances microtubule growth to pattern the axon for synaptogenesis and guide presynaptic cargo distribution.

## RESULTS

### Axonal presynaptic accumulation sites along human i^3^Neuron axons are hotspots for anterograde and retrograde SVP pauses and microtubule polymerization events

In primary rat hippocampal neurons, local enrichment of microtubule polymerization influences SVP pausing and retention at *en passant* presynaptic sites [7]. Here, we confirm that this phenomenon is conserved in human neurons using a genetically engineered iPSC line with a doxycycline-inducible Neurogenin-2 (NGN2) expression cassette inserted into the AAVS1 safe harbor locus to generate a homogenous population of glutamatergic cortical-like neurons termed i^3^Neurons [24, 25]. We imaged i^3^Neurons expressing fluorescently labeled synaptophysin (mScarlet-Syp), a synaptic vesicle transmembrane protein, at DIV35 to track axonal SVP movement. We observed robust SVP flux in human i^3^Neurons in both the anterograde and retrograde directions, punctuated by discrete pausing and retention events (**Figure 1A-C**), similar to previous observations in rat hippocampal neurons [7]. Anterograde SVP flux was observed at a rate of 3.2 vesicles/minute moving at an average instantaneous velocity of 2.7 µm/s in i^3^Neurons (**Figure 1C, S1A**). In the retrograde direction, SVP flux was observed at 0.9 vesicles/minute, with SVPs moving at 2.2 µm/s (**Figure 1C, S1A**).

**Figure 1.**
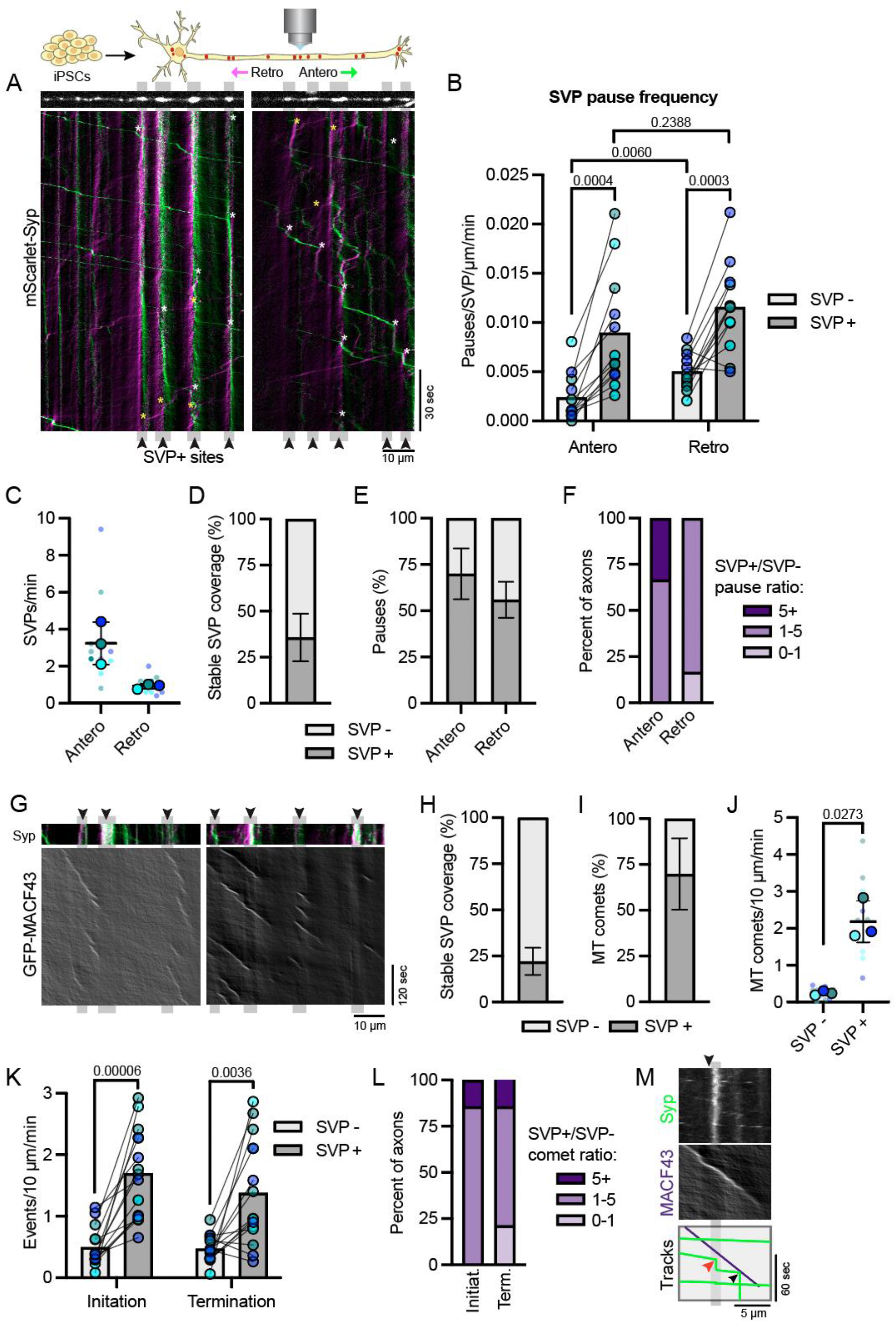
Presynaptic accumulation sites along human i^3^Neuron axons are hotspots for anterograde and retrograde SVP pauses and enriched for microtubule polymerization. **(A)** Examples of axonal motility of SVPs in DIV35 i^3^Neurons expressing mScarlet-Syp. Anterograde and retrograde tracks are green and magenta, respectively. Upper panels: still image of axonal mScarlet-Syp signal. Lower panels: kymograph of SVPs imaged at 200 ms per frame. SVP+ sites of stably accumulated SVPs are emphasized in gray. Examples of anterograde and retrograde SVP pauses are highlighted with white and yellow asterisks, respectively. **(B)** Anterograde and retrograde SVP pause events standardized for vesicle flux and distance in axonal areas lacking stable SVPs (SVP-, light gray) and populated by stable SVPs (SVP+, dark gray). Each of the paired data points represent SVP- and SVP+ pausing frequency within one axon. Reported *p*-values are from multiple paired t-test (between SVP+ and SVP-values) and from one-way repeated measures ANOVA (between anterograde and retrograde values). **(C)** SVP flux (vesicles/min) in the anterograde and retrograde direction. Error bars represent the standard deviation of the replicate means. **(D)** Ratios of SVP pause frequency in SVP+/SVP-regions. Percent of axons with ratios of 0-1, 1-5, and 5+ are displayed in light, medium, and dark purple. 100% of axons exhibit an increase in anterograde pause frequency at SVP+ sites (SVP+/SVP-ratio ranging from 1.9-21), and 83% of axons exhibit an increase in retrograde pause frequency at SVP+ sites (ratio ranging from 0.7-4.8). **(E)** Stable SVP coverage as percent of total axon distance at DIV35 (stable SVP µm/total analyzed axon µm, dark gray). Error bars represent the standard deviation of the data set. **(F)** Percent of SVP pauses that occur at stable SVP regions (SVP+ pauses/total pauses, dark gray) in the anterograde and retrograde directions. Error bars represent the standard deviation of the data set. **(G)** Representative kymographs of axonal microtubule comet events imaged 1 frame per second in DIV21 i^3^Neurons expressing MACF43-GFP and mScarlet-Syp. Upper panels: mScarlet-Syp signal reveals SVP+ sites. Lower panels: GFP-MACF43 kymographs display location and timing of new microtubule polymerization. SVP+ sites are emphasized in gray. **(H)** Stable SVP coverage as percent of total axon distance at DIV21 (stable SVP µm/total analyzed axon µm, dark gray). Error bars represent the standard deviation of the data set. **(I)** Percent of microtubule (MT) comets that occur at stable SVP regions (SVP+-associated comets/total comets, dark gray). Error bars represent the standard deviation of the data set. **(J)** Microtubule comet events standardized to analyzed kymograph distance and time. SVP+ regions exhibit an 8.8-fold increase in microtubule growth events, determined by counting comets that initiate, terminate, or pass through SVP+ sites. **(K)** Microtubule comet initiation and termination events standardized to distance and time in axonal areas lacking stable SVP (SVP-, light gray) and populated by stable SVP (SVP+, dark gray). **(L)** Ratios of microtubule comet initiation and termination in SVP+/SVP-regions. Percent of axons with ratios of 0-1, 1-5, and 5+ are displayed in light, medium, and dark purple. 100% of axons exhibit an increase in microtubule comet initiation at SVP+ sites (ratios ranging from 1.4-35), and 79% of axons exhibit an increase in microtubule comet termination at SVP+ sites (ratios ranging from 0.6-48). **(M)** Representative kymograph of SVP movement (left panel, mScarlet-Syp) and microtubule polymerization (middle panel, GFP-MACF43). Right panel provides merged position of motile SVP tracks (green), stationary SVP+ sites (gray), and microtubule comet track (purple) showing SVP pause at a SVP+ site (red arrowhead) and at plus end of microtubule (black arrow).

We found that i^3^Neuron axons are punctuated by sites of stable accumulation of presynaptic components, or SVP+ regions. SVPs preferentially paused at these sites in both the anterograde and retrograde direction (**Figure 1A, B**). Despite stable SVP accumulations occupying only 34% of the total analyzed axon distance (**Figure 1D**), 70% of anterograde and 56% of retrograde SVPs paused specifically at these sites (**Figure 1E**). All axons exhibited increased anterograde SVP pausing at SVP+ sites compared to regions lacking stable SVPs, with 67% showing an 1-5-fold increase and 33% showing a 5+-fold increase (**Figure 1B, F**). In the retrograde direction localized SVP pausing was also observed, with 83% of axons exhibiting a 1-5-fold increase in SVP pausing at SVP+ regions (**Figure 1B, F**). In contrast, in primary rat neurons only anterograde-directed SVPs demonstrate a preference for pausing at stable SVP accumulation sites [7]. Our results demonstrate that both anterograde and retrograde SVPs preferentially pause at SVP+ regions in human i^3^Neurons, with a particularly strong preference observed for SVP+ pausing in the anterograde direction. Thus, while there are differences observed between rat and human cultured neurons, the pausing preference for anterograde-moving SVPs at stable SVP accumulations is conserved across systems.

To interrogate whether SVP+ sites are also hotspots for microtubule polymerization events, we transiently expressed the microtubule plus-end marker GFP-MACF43 and the presynaptic cargo mScarlet-Syp in i^3^Neurons and imaged each channel at 1 frame per second to observe microtubule growth events, or “comets”, in relation to SVP+ sites. MACF43 acts as a microtubule +TIP via interaction with endogenous end-binding proteins (EBs) [26]. As no significant changes to SVP pausing behavior were detected between DIV21 and DIV35 (Figure 1B and S1B), all subsequent experiments were performed at DIV21 to facilitate data collection. Microtubule polymerization events are strikingly enriched at SVP+ sites (**Figure 1G**). While only 22% of the total analyzed axon is occupied by stable SVPs at DIV21 (**Figure 1H**), 70% of microtubule comets initiate, pass through, or terminate at these SVP+ regions (**Figure 1I**). When standardized for distance and time, SVP+ sites exhibited an 8.8-fold increase in comet events (comets that initiate, terminate, or pass through the site) compared to regions lacking accumulated presynaptic cargos (**Figure 1J**). SVP+ sites are hotspots for both comet initiation and termination events (**Figure 1K, L**), demonstrating that microtubule polymerization preferentially starts and stops within these regions. Microtubule plus ends enriched for GTP-tubulin promote KIF1A detachment [7], and we accordingly observed SVP pauses initiating at the plus end of microtubules (black arrow, **Figure 1M**) and SVP+ sites (red arrowhead, **Figure 1M**). Our data reveal that i^3^Neurons, like primary rat hippocampal neurons [7], exhibit preferential microtubule polymerization and anterograde SVP pauses at axonal regions occupied by stable SVPs.

### Spastin disruption alters density of axonal microtubule polymerization events

To examine whether spastin regulates axonal microtubule polymerization, we used CRISPR interference (CRISPRi; **Figure 2A**) to knockdown spastin protein to 18% of its wild-type level (**Figure S2A**). We compared these spastin knockdown i^3^Neurons (Sp KD) to non-targeting CRISPRi i^3^Neurons (Control) and to i^3^Neurons transiently overexpressing SNAP-tagged spastin (Sp OE). Two distinct isoforms of spastin are expressed from different initiation codons: M1 and M87. We chose to overexpress the M87 isoform as it is the predominant isoform in the developing CNS [27]. SVPs contain numerous membrane-bound cargos destined for the synapse, including synaptophysin-1 (Syp) and synaptobrevin-2 (Syb) [28]. For ease of co-visualization of both presynaptic cargo and microtubule comets, we generated a bicistronic vector that simultaneously expresses mScarlet-Syb and GFP-MACF43. We noted very similar pausing behavior at SVP+ sites using distinct SVP markers (Syp and Syb) in i^3^Neurons at both DIV21 and DIV35, revealing consistency across experimental paradigms.

**Figure 2.**
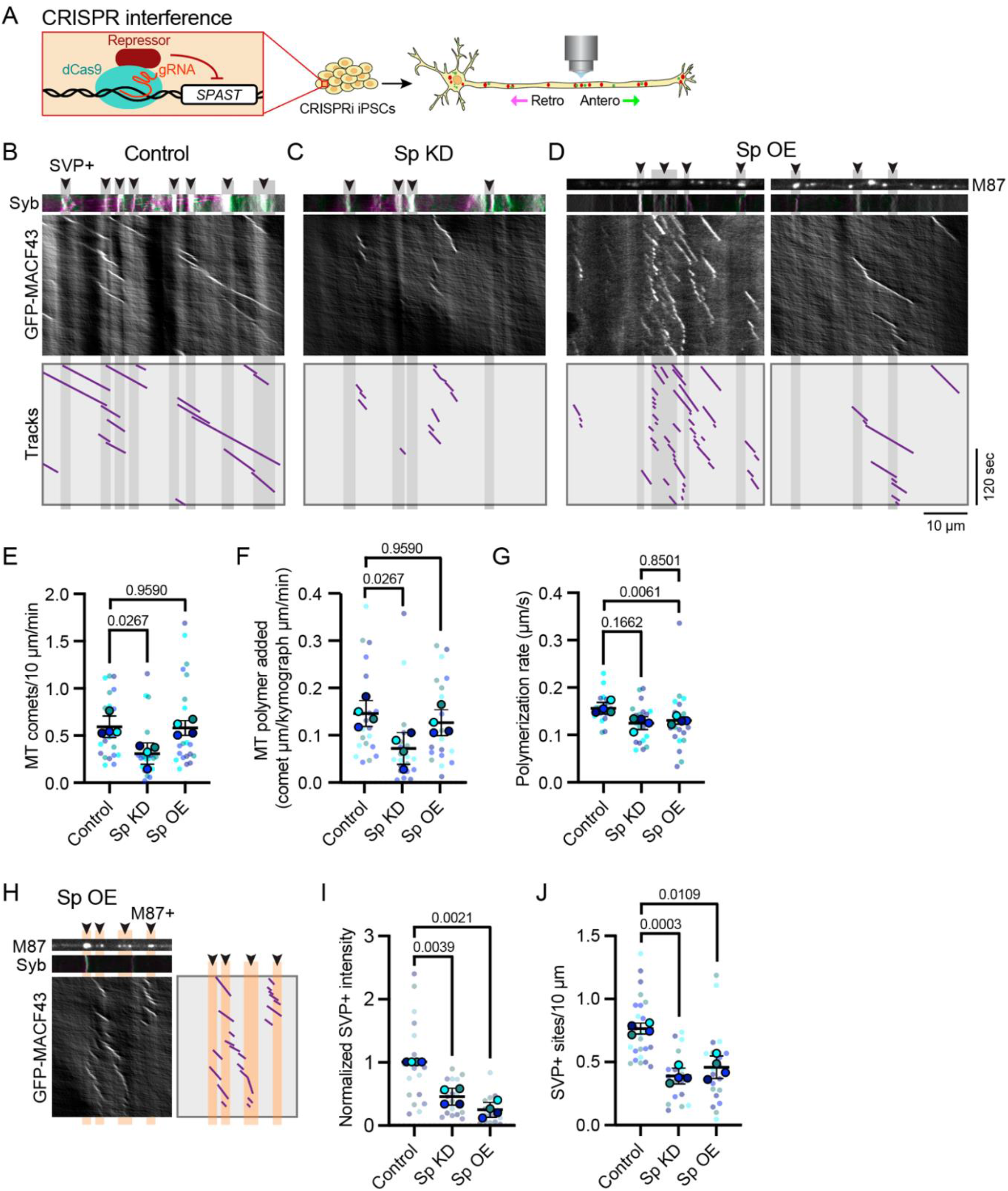
Spastin disruption alters density of axonal microtubule polymerization events and organization of presynaptic accumulations. **(A)** Schematic of CRIPSR-interference (CRISPRi) regulation of gene expression. Guide RNA (gRNA) targets nuclease-deficient dCas9 fused to a transcription repressor to the gene of interest, reducing target gene transcription. **(B-D)** Representative kymographs of axonal microtubule comet events in DIV21 i^3^Neurons expressing bicistronic mScarlet-Syb-IRES-MACF43-GFP for CRISPRi non-targeting control (Control; **B**), spastin CRISPRi-mediated knockdown (Sp KD; **C**), and spastin M87 isoform transient overexpression (Sp OE; **D**). Upper panels: mScarlet-Syb (Syb) signal demarks SVP+ sites (**B-D**) and SNAP-M87 reveals overexpressed spastin localization (M87, top panel of **D**). Middle panels: GFP-MACF43 kymographs display location and timing of new microtubule polymerization. Lower panels: Microtubule comet tracks are provided in purple. SVP+ sites are emphasized in gray. **(E)** Microtubule comet density (comets/10 µm of axon/min) for Control, Sp KD, and Sp OE. Reported *p*-values are from Friedman’s non-parametric test for multiple comparisons, as the data failed the Shapiro-Wilk test for normality. **(F)** New microtubule polymer added (comet µm/analyzed axon µm/min) for Control, Sp KD, and Sp OE. Reported *p*-values are from Friedman’s non-parametric test for multiple comparisons, as the data failed the Shapiro-Wilk test for normality. **(G)** Microtubule polymerization rate (µm/second) from Control, Sp KD, and Sp OE. Reported *p*-values are from one-way repeated measures ANOVA. **(H)** Representative kymograph of Sp OE axon highlighting location of stable spastin M87 in relation to microtubule comets. Top left panel: still image of SNAP-M87 signal. Middle left panel: Syb signal reveals one SVP+ site. Bottom left panel: GFP-MACF43 kymograph reveals location and timing of new microtubule polymerization. Right panel: Purple tracks corresponding to microtubule comets. Stable M87 position is emphasized in orange. **(I)** Normalized Syp intensity at SVP+ sites Reported *p*-values are from one-way repeated measures ANOVA. **(J)** SVP+ site density (number of Stable SVP+ puncta/10 µm; **J**) for Control, Sp KD, and Sp OE. Reported *p*-values are from one-way repeated measures ANOVA. For all plots, errors bars represent the standard deviation of the replicate means.

Microtubule comet analysis reveals that spastin KD leads to significantly fewer axonal microtubule comet events (**Figure 2B, C, E**) and less total microtubule polymer added (sum of comet lengths standardized to kymograph distance and time; **Figure 2B, C, F**) compared to control i^3^Neurons. In contrast, spastin OE did not significantly alter the number of microtubule comet events (**Figure 2B, D, E**) or the total microtubule polymer added (**Figure 2B, D, F**). However, consistent with spastin’s potential to act as a microtubule network amplifier or dismantler [11–13, 23], axons overexpressing spastin exhibited more variability, with some axon segments displaying numerous microtubule comet events (**Figure 2D**, left panel), while others displayed few, or no, microtubule comet events (**Figure 2D**, right panel; **Figure S2B**). Interestingly, either decreasing or increasing neuronal spastin levels caused similar shifts in microtubule polymerization rate, with a significant reduction observed in spastin overexpression and a trend toward shorter polymerization in spastin knockdown (**Figure 2G**). Both knockdown and overexpression of spastin induced a trend toward shorter polymerization distance and longer duration (**Figure S2C, D**). Visualizing SNAP-M87 spastin revealed that 60% of microtubule comets events corresponded to stationary spastin puncta (**Figure 2H; Figure S2E**), despite spastin occupying only 14% of the total analyzed axon distance (**Figure S2F**). Taken together, these data indicate that spastin acts within the axon to amplify new microtubule polymerization events.

Both spastin KD and OE axons exhibited a significant decrease in both intensity and number of SVP+ sites (**Figure 2I, J**), indicating that stable SVP patterning and presynaptic cargo accumulation are disrupted in parallel upon modulation of spastin levels. In wild-type neurons, the amount of new microtubule polymer added correlated with SVP+ coverage of the axonal region (**Figure S2G**). This relationship is disrupted in both the spastin KD and OE conditions, indicating that precise regulation of microtubule polymerization via spastin influences presynaptic patterning.

### Spastin disruption interrupts the positioning of new microtubule polymerization events and SVP+ organization

As spastin levels influence the density of axonal microtubule comets and SVP accumulation, we next asked whether the positioning of microtubule polymerization is disrupted upon spastin KD or OE. We analyzed kymographs generated from control, spastin KD, or spastin OE i^3^Neurons expressing the mScarlet-Syb-IRES-GFP-MACF43 bicistronic vector and analyzed the position of the microtubule comets in relation to stable SVP+ sites (**Figure 3A-C**). SVP+ comet events (shown in pink) were categorized as MACF43 comets that initiate, pass through, or terminate directly within stable SVP+ regions, with SVP-comets, i.e. comets not associated with stable SVP sties, in blue. In control axons, microtubule comets were enriched in regions inhabited by stable SVP accumulations; this can be appreciated by the increased microtubule comet density in the vicinity of SVP+ sites (emphasized in gray), as well as the increased ratio of pink SVP+ tracks to blue SVP-tracks in the example kymograph tracing (**Figure 3A**). The wild-type results presented in Figure 1 were echoed in CRISPRi non-targeting control axons, with increased microtubule comet initiation and termination observed at SVP+ sites (**Figure 1K, L and 3D, E**). 100% of axons examined exhibited an increased ratio in SVP+/SVP-comet initiation events (ratios ranging from 1.2-15) and 86% of axons exhibited increased termination event ratios (ratios ranging from 0.5-20; **Figure 3E**). In contrast, CRISPRi-mediated spastin KD led to a striking reduction in SVP+ microtubule comets (**Figure 3B**). While regions lacking stable SVP accumulations (SVP-) exhibited no change in microtubule comet density between control and spastin KD conditions, SVP+ regions demonstrated a significant decrease in associated microtubule polymerization events when spastin was depleted (**Figure 3F**). Spastin KD decreased the preference for both comet initiation and termination in SVP+ regions (**Figure 3D**), shifting the balance to fewer axons demonstrating enrichment of microtubule polymerization events at SVP+ sites (**Figure 3E**). Spastin OE disrupted microtubule comet positioning in the opposite way: SVP-regions exhibited a significant increase in microtubule comet density while SVP+ regions exhibited no change when spastin is overexpressed (**Figure 3C-F**). In spastin OE axons, there appears to be a “spill-over” effect where mislocalized comets are found outside of SVP+ sites, often in immediately adjacent regions. Accordingly, we observe ectopically expressed SNAP-M87 spastin both near and between SVP+ sites (**Figure 2H, 3C**). Taken together, our live-imaging results indicate that spastin influences axonal microtubule comet density and position, with spastin KD leading to fewer microtubule polymerization events within SVP+ sites and spastin OE leading to more polymerization events outside SVP+ regions.

**Figure 3.**
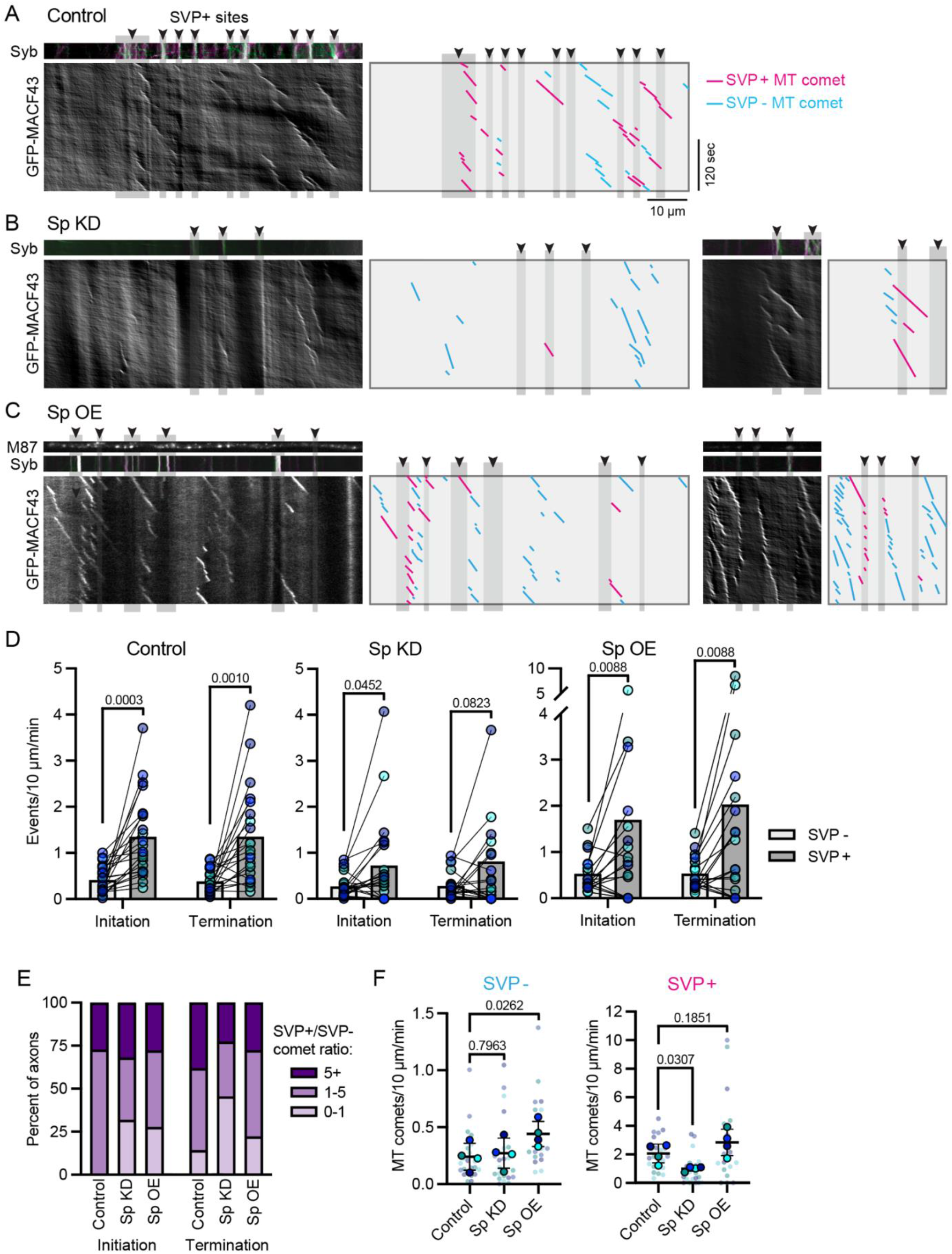
Spastin disruption interrupts positioning of new microtubule polymerization events. **(A-C)** Representative kymographs of axonal microtubule comet events in DIV21 i^3^Neurons expressing mScarlet-Syb-IRES-MACF43-GFP for CRISPRi non-targeting control (Control; **A**), spastin CRISPRi-mediated knockdown (Sp KD; **B**), and spastin M87 isoform transient overexpression (Sp OE; **C**). Upper panels: mScarlet-Syb (Syb) signal demarks SVP+ sites (**A-C**) and SNAP-M87 (M87, top panels of **C**) displays overexpressed spastin localization. Lower panels: GFP-MACF43 kymographs and tracks reveal location and timing of new microtubule polymerization. SVP+ sites are emphasized in gray. Corresponding tracks differentiate microtubule comets not associated with SVP+ regions (SVP-, blue) from comets that that initiate, terminate, and/or pass through a SVP+ site (SVP+, pink). **(D)** Microtubule initiation and termination events (events/10 µm/min) in axonal areas lacking stable Syb (SVP-, light gray) and populated by stable Syb (SVP+, dark gray) for Control, Sp KD, and Sp OE. Reported *p*-values are from multiple paired t-test. **(E)** Ratios of microtubule comet initiation and termination in SVP+/SVP-regions. Percent of axons with ratios of 0-1, 1-5, and 5+ are displayed in light, medium, and dark purple. 100% of control axons exhibit an increased microtubule comet initiation frequency at SVP+ sites. Both Sp KD and Sp OE shifts the SVP+/SVP-ratio to more comets initiating outside of SVP+ regions. 86% of control axons exhibit an increased microtubule comet termination frequency at SVP+ sites. Sp KD alters the SVP+/SVP-ratio of termination events so only 54% of axons exhibit a ratio higher than 1 (i.e. pauses enriched at SVP+ sites). **(F)** Microtubule comet density in SVP-(left graph) or SVP+ (right graph) regions in Control, Sp KD, and Sp OE axons. Sp OE leads to more microtubule comets positioned outside SVP+ regions, while Sp KD leads to fewer SVP+ microtubule comets. Errors bars represent the standard deviation of the replicate means. Reported *p*-values are from one-way repeated measures ANOVA.

### Spastin-deficient axons exhibit a decrease in overall anterograde SVP pause and retention events

To directly test whether spastin levels influences SVP trafficking behavior, we rapidly live imaged mScarlet-Syb in CRISPRi spastin KD and non-targeting control axons at 200 ms/frame (**Figure 4A**). In general, spastin KD does not grossly alter SVP flux (no change in anterograde flux, trend toward decreased retrograde flux; **Figure 4B**). SVP velocity also does not significantly change upon spastin KD (**Figure 4C**). However, spastin depletion caused a marked reduction in SVP pause frequency, the percent of SVPs that exhibit a pause, and SVP retention, specifically in the anterograde direction (**Figure 4D-F**). SVP pause duration, in contrast, is unaffected (**Figure 4G**), suggesting that when an SVP pauses, it reengages with the microtubule network similarly in both control and spastin KD conditions to initiate its next processive run.

**Figure 4.**
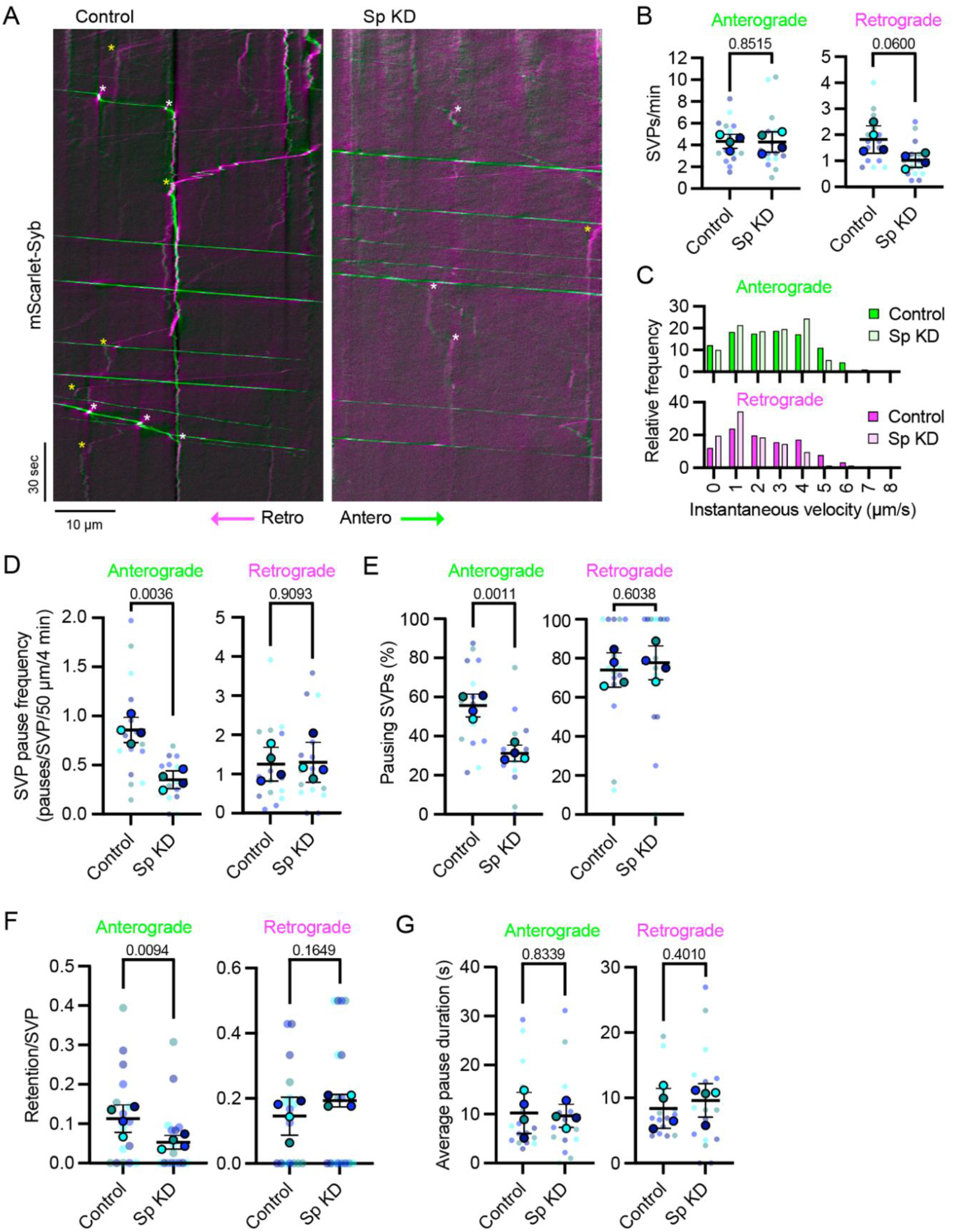
Spastin deficient axons exhibit a decrease in overall anterograde SVP pause events. **(A)** Representative kymographs of axonal SVP movement in DIV21 i^3^Neurons expressing mScarlet-Syb for CRISPRi non-targeting control (Control) and spastin CRISPRi-mediated knockdown (Sp KD). Examples of anterograde and retrograde SVP pauses/retentions are highlighted with white and yellow asterisks, respectively. **(B)** SVP flux (vesicles/min) for Control and Sp KD in the anterograde and retrograde direction. For plots **B, D-F**, errors bars represent the standard deviation of the replicate means and reported *p*-values are from a t-test comparing the means of four experimental replicates. **(C)** Histograms of instantaneous velocities in µm/second for anterograde (upper) and retrograde (lower) Control and Sp KD SVPs. SVP average instantaneous velocities were not significantly different between Control and Sp KD in the anterograde (*p=*0.8619) or retrograde (*p=*0.5860) direction, as determined by a t-test comparing four experimental replicate means. **(D)** Total anterograde (left) and retrograde (right) SVP pause frequency standardized to vesicle flux and kymograph distance for Control and Sp KD. **(E)** Percent of pausing SVPs (SVPs that experience pause/total SVPs) for Control and Sp KD in the anterograde (left) and retrograde (right) directions. **(F)** Retention events per SVP for anterograde (left) and retrograde (right) SVPs in Control and Sp KD axons. **(G)** Average pause duration in seconds for Control and Sp KD SVPs in the anterograde (left) and retrograde (right) directions.

### Depletion of neuronal spastin interrupts localization of anterograde presynaptic cargo pausing and retention

We next asked whether the observed changes in SVP pausing behavior upon spastin depletion can be ascribed to transport delivery defects at SVP+ sites. Kymographs generated from rapid live imaging of control or spastin KD i^3^Neurons expressing mScarlet-Syb were analyzed for SVP pauses and retention in relation to stable SVP+ accumulations (**Figure 5A**). Although we analyzed axonal regions with similar SVP+ coverage (18% and 15% of control and spastin KD axons, respectively), anterograde pauses at SVP+ regions decreased from 52% in control to 25% when spastin is depleted (**Figure 5B, C**). In agreement with wild-type i^3^Neuron data presented in Figures 1 and S1, all DIV21 control axons exhibited increased anterograde SVP pausing at SVP+ sites compared to regions lacking stable SVPs, with 69% showing an 1-5-fold increase and 31% showing a 5+-fold increase (**Figure 5D, E**). Spastin KD axons exhibited a much less pronounced enrichment of SVP pauses at SVP+ sites (**Figure 5D**) and decreased the SVP+/SVP-pause ratio (**Figure 5E**), reminiscent of the shift observed for SVP+/SVP-microtubule comets (**Figure 3D, E**). SVP retention at SVP+ sites was similarly decreased upon spastin KD (**Figure 5F, G**). Indeed, anterograde SVP retentions shifted from nearly all occurring at SVP+ regions in control axons to less than a third in spastin KD axons (96% vs. 30%, respectively; **Figure 5G**).

**Figure 5.**
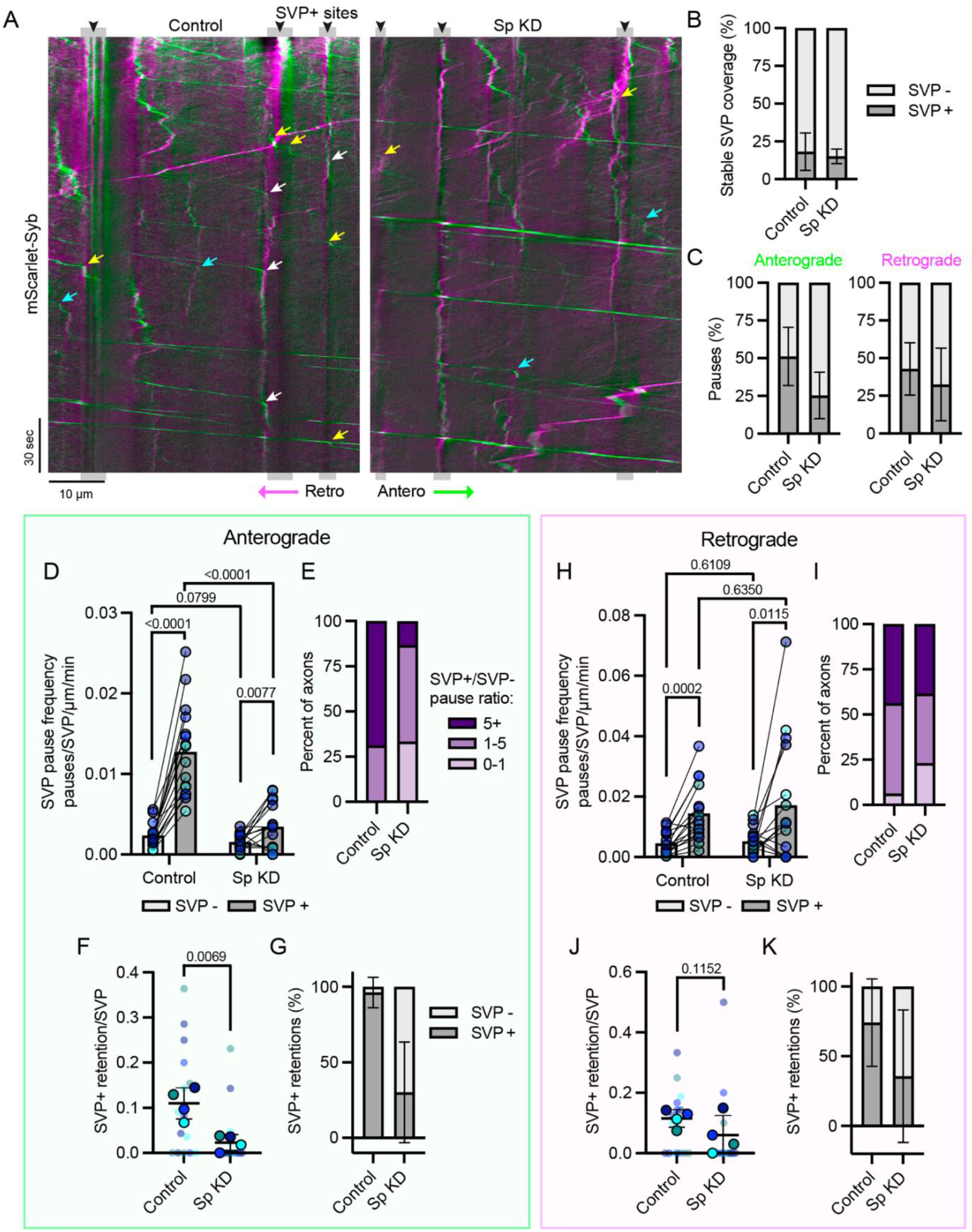
Depletion of neuronal spastin interrupts localization of anterograde presynaptic cargo pausing and retention. **(A)** Representative kymographs of axonal SVP movement in DIV21 i^3^Neurons expressing mScarlet-Syb for CRISPRi non-targeting control (Control) and spastin CRISPRi-mediated knockdown (Sp KD). Anterograde and retrograde movement is displayed in green and magenta, respectively. SVP+ sites are emphasized in gray. Arrows highlight examples of anterograde SVP retention (white), pausing at stable SVP+ sites (yellow), and pausing at regions lacking stable SVPs (blue). **(B)** Axonal SVP+ coverage as percent of total axon distance (stable SVP µm/total analyzed axon µm, dark gray) for Control and Sp KD. Error bars represent the standard deviation of the data set. **(C)** Percent of SVP pauses that occur at SVP+ (dark gray) or SVP-(light gray) regions in the anterograde (left) or retrograde (right) directions. Error bars represent the standard deviation of the data set. **(D)** Anterograde SVP pause events standardized for vesicle flux and distance in axonal areas lacking stable SVPs (SVP-, light gray) and populated by stable SVPs (SVP+, dark gray) for Control and Sp KD axons. Paired data points represent SVP-and SVP+ pausing frequency within one axon. Reported *p*-values are from multiple paired t-test (between SVP+ and SVP-values) and from one-way repeated measures ANOVA (between Control and Sp KD values). **(E)** Ratios of SVP pause frequency in SVP+/SVP-regions in the anterograde direction. Percent of axons with ratios of 0-1, 1-5, and 5+ are displayed in light, medium, and dark purple. **(F)** Number of retentions at SVP+ sites per anterograde SVP for Control and Sp KD. Sp KD significantly decreases the number of SVP+ retentions. Error bars represent the standard deviation of the replicate means and provided *p*-values were determined by t-test comparing the four experimental replicate means. **(G)** Percent of anterograde SVPs retained at SVP+ regions (SVP+ retentions/total retentions; dark gray) for Control and Sp KD. **(H)** Retrograde SVP pause events standardized for vesicle flux and distance in axonal areas lacking stable SVPs (SVP-, light gray) and populated by stable SVPs (SVP+, dark gray) for Control and Sp KD axons. Reported *p*-values for SVP+ and SVP-comparisons were determined by multiple paired t-test. *p*-values for Control and Sp KD comparisons were determined by one-way repeated measures ANOVA. **(I)** Ratios of SVP pause frequency in SVP+/SVP-regions in the retrograde direction. Percent of axons with ratios of 0-1, 1-5, and 5+ are displayed in light, medium, and dark purple. **(J)** Number of retentions at SVP+ sites per retrograde SVP for Control and Sp KD. Error bars represent the standard deviation of the replicate means and provided *p*-values were determined by t-test comparing the four experimental replicate means. **(K)** Percent of retrograde SVPs retained at SVP+ regions (SVP+ retentions/total retentions; dark gray) for Control and Sp KD.

The consequences of spastin depletion on retrograde SVP trafficking were much less dramatic (**Figure 5H-K**). Both control and spastin KD conditions exhibited increased retrograde pause frequency at SVP+ sites (**Figure 5H**). However, spastin KD caused a subtle shift toward fewer axons experiencing an enrichment in SVP+/SVP-retrograde pause events (**Figure 5I**), a trend toward fewer SVP+ retentions per SVP (**Figure 5J**), and a decrease in the percent of retrograde retentions occurring at SVP+ sites from 74% in control to 36% in spastin KD (**Figure 5K**). These data highlight a previously unappreciated role for spastin in regulating presynaptic cargo trafficking and delivery to specific sites along the axon, with strong defects observed in anterograde SVP pausing/retention and less pronounced changes in retrograde trafficking upon spastin depletion.

### Presynaptic accumulations and *bona fide* presynapses are capable of cycling synaptic vesicles and are enriched for endogenous spastin

i^3^Neurons have previously been employed to generate and study human neurons in culture [25, 29, 30], but synapse formation has not yet been thoroughly characterized in this system. We discovered that for i^3^Neurons in monoculture at DIV21, a commonly used end-point for i^3^Neuron experiments, most presynaptic puncta are not apposed by postsynaptic counterparts as revealed by Synapsin I/II (Syn), PSD-95, and MAP2 immunocytochemistry (**Figure 6A, B**). In fact, of the Syn puncta determined to be the correct size and intensity to be potential presynapses in i^3^Neuron monocultures (using FIJI SynapseJ macro [31] and 3D Objects Counter), only 15% at DIV14 and 18% at DIV21 co-localized with PSD-95 along MAP2+ dendrites. Overall, we noted a sparse 0.9 and 1.2 synapses per 100 µm^2^ of somatodendritic area at DIV14 and DIV21, respectively (**Figure 6B, C**). Thus, it is likely that only a minority of the stable presynaptic accumulations characterized by live imaging in Figures 1-5 are apposed by postsynapses. Instead, they may reflect a presynaptic structure upstream of synaptogenesis, which we term ‘protosynapse’. In support of this idea, i^3^Neurons co-cultured with primary rat astrocytes reach more mature synaptic states (**Figure 6A-C**). While astrocyte co-culture is not sufficient to significantly increase i^3^Neuron synapse density at DIV21, DIV42 i^3^Neurons co-cultured with astrocytes exhibited a dramatic increase in both percentage of Syn puncta apposed by postsynaptic marker PSD95 (**Figure 6B**) as well as synapse density per area (**Figure 6C**). By DIV42, i^3^Neurons exhibited 39% of Syn puncta co-localized with PSD-95 along MAP2+ dendrites and an average of 4.4 synapses per 100 µm^2^ of somatodendritic area. This immunocytochemical characterization of synapses in DIV42 i^3^Neurons co-cultured with astrocytes is remarkably similar to what has been observed in primary rat neuron cultures at DIV14, where approximately 40% of synaptophysin puncta colocalize with postsynaptic marker AMPA-R, leading to 5 synapses per 100 µm^2^ [32]. Primary rat cultures also exhibit a shift over time in percent of presynaptic puncta apposed by postsynaptic markers [32], suggesting that presynaptic puncta transition to become apposed by post-synaptic partners in both murine and human neuron models.

**Figure 6.**
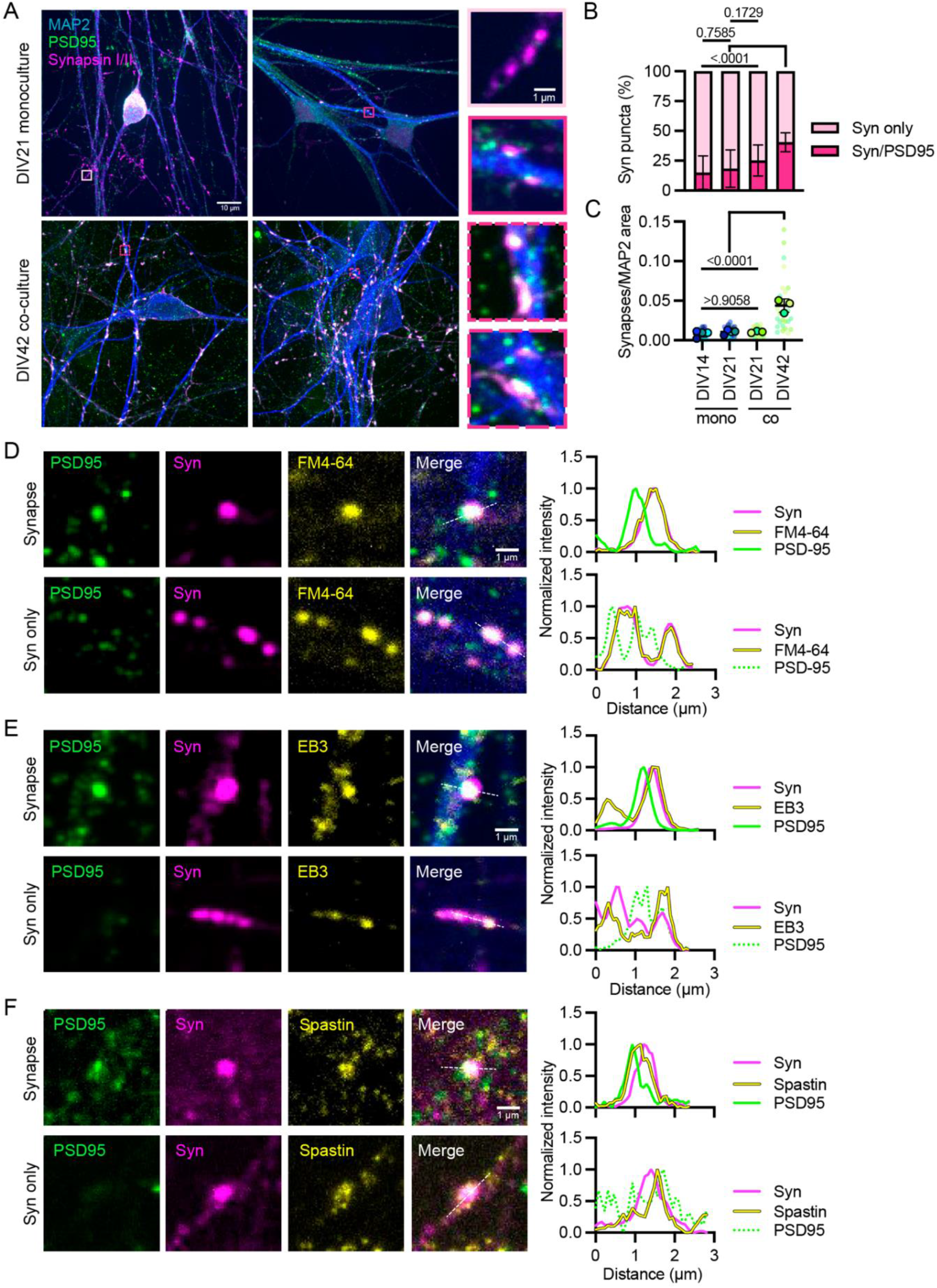
Spastin is enriched at both synapses and protosynapses in i^3^Neurons. **(A)** Representative immunocytochemistry MAX-projection images of endogenous MAP2 (somatodendritic compartment, blue), PSD95 (postsynaptic marker, green), and Synapsin I/II (Syn, presynaptic marker, magenta) in DIV21 i^3^Neurons in monoculture (top) and DIV42 i^3^Neurons co-cultured with primary rat astrocytes (bottom). Light pink box/inset highlights presynapse puncta without postsynaptic counterpart (i.e. protosynapse). Dark pink solid and dashed boxes/insets highlight *bona fide* synapses with apposed pre- and postsynapse. **(B)** Percent of Syn puncta that are apposed by PSD95 (dark pink) or alone (light pink) in i^3^Neurons at DIV14 and DIV21 in monoculture (mono) and i^3^Neurons at DIV21 and DIV42 co-cultured (co) with primary rat astrocytes. Error bars represent the standard deviation of the data set and reported *p*-values are from one-way repeated measures ANOVA. **(C)** Synapses per µm^2^ of MAP2 somatodendritic area for monoculture (mono) i^3^Neurons at DIV14 and DIV21 and co-cultured (co) i^3^Neurons with primary rat astrocytes at DIV21 and DIV42. Pre- and post-synaptic puncta were determined using FIJI SynapseJ macro and 3D Object Counter on z-stack images. Error bars represent the standard deviation of the replicate means and reported *p*-values are from one-way repeated measures ANOVA. **(D-F)** Max intensity projections of *bona fide* synapses (top panels) and Syn only protosynapses (bottom panels) in immunostained DIV21 monoculture wild-type i^3^Neurons. White dashed line demarks region used for intensity profile of each channel provided to the right. **(D)** Syn+ presynaptic puncta (magenta) of *bona fide* synapses (top) and Syn only protosynapses (bottom) contain cycling synaptic vesicles as visualized by FM4-64 dye (yellow). **(E)** +TIP EB3 (yellow) reveals microtube plus-ends at Syn+ (magenta) *bona fide* synapses (top) and Syn only protosynapses (bottom). **(F)** Spastin (yellow) is found at Syn+ (magenta) *bona fide* synapses (top) and Syn only protosynapses (bottom).

We sought to characterize both the *bona fide* synapses consisting of pre- and postsynaptic densities as well as protosynapses consisting of presynaptic marker alone in DIV21 monoculture i^3^Neurons, which reflect the synaptic state of the i^3^Neurons used for live imaging. When synapses are present in i^3^Neurons, they contain cycling vesicle clusters as determined by the uptake of FM4-64 dye upon K+ stimulation (**Figure 6D**, upper panels). Protosynaptic (Syn only) puncta within i^3^Neuron axons are also capable of cycling synaptic vesicles independent of postsynaptic contacts (**Figure 6D**, lower panels), in agreement with data generated from primary rat neurons [5].

Endogenous microtubule +TIP EB3 is found associated with both *bona fide* presynapses and protosynapses (**Figure 6E**). EB3 puncta correspond to newly polymerized microtubule plus ends; these immunofluorescence results corroborate the observed enrichment of microtubule comets at SVP+ sites upon i^3^Neuron live imaging (**Figure 1G, 2B, 3A**). Immunocytochemistry of endogenous spastin reveals enrichment at both *bona fide* synapses and protosynapses (**Figure 6F**), where it is ideally positioned to locally regulate microtubule dynamics at presynaptic accumulations punctuating the axon. Within *bona fide* synapses, spastin appears to be positioned at both the pre- and postsynapse. Spastin co-localization at the postsynapse is consistent with previous reports of spastin knockout leading to dendritic spine morphology and density defects in mice [21, 33].

### Human heterologous synapse assay models presynapse formation containing cycling synaptic vesicles and spastin

With spastin potentially regulating both pre- and postsynaptic compartments, we sought to specifically interrogate spastin’s role at the presynapse using an innovative human heterologous synapse model. In this system, non-neuronal human embryonic kidney (HEK) cells expressing Neuroligin-1 (NL1) are introduced to microfluidically isolated i^3^Neuron axons, which recognize and form presynaptic connections with the presented postsynaptic ligand within 24 hours. Where axons cross non-NL1 expressing HEK cells (example outlined in white in **Figure 7A**), no presynapses are established, but where axons encounter HEK cells expressing NL1, robust presynaptic connections are formed (**Figure 7A**). These heterologous presynapses are enriched in presynaptic markers Syn, Syp, Syb, and the excitatory transmitter VGLUT1, and contain synaptic vesicles that can cycle upon depolarization, as shown by FM4-64 dye uptake (**Figure 7A-D**). Interestingly, the density of heterologous synapses in axons crossing NL1+ HEK cells is similar to the density of SVP+ sites, determined using live imaging data, and protosynapses, determined by immunocytochemistry in microfluidically isolated axons, with approximately 1 puncta per 10 µm of axon in all instances (**Figure S3A**). Similar to *bona fide* presynapses and protosynapses, heterologous presynapses are also enriched for the +TIP EB3 and spastin (**Figure 7E, F**; white insets). They do not, however, contain the microtubule nucleator γ-tubulin (**Figure 7E**, white inset), which had previously been implicated in presynaptic microtubule regulation upon stimulation in rodent neurons [8]. Our immunofluorescence results indicate functional consistency across protosynapses, *bona fide* presynapses, and heterologous presynapses, with all capable of cycling synaptic vesicles and enriched for spastin and microtubule plus ends.

**Figure 7.**
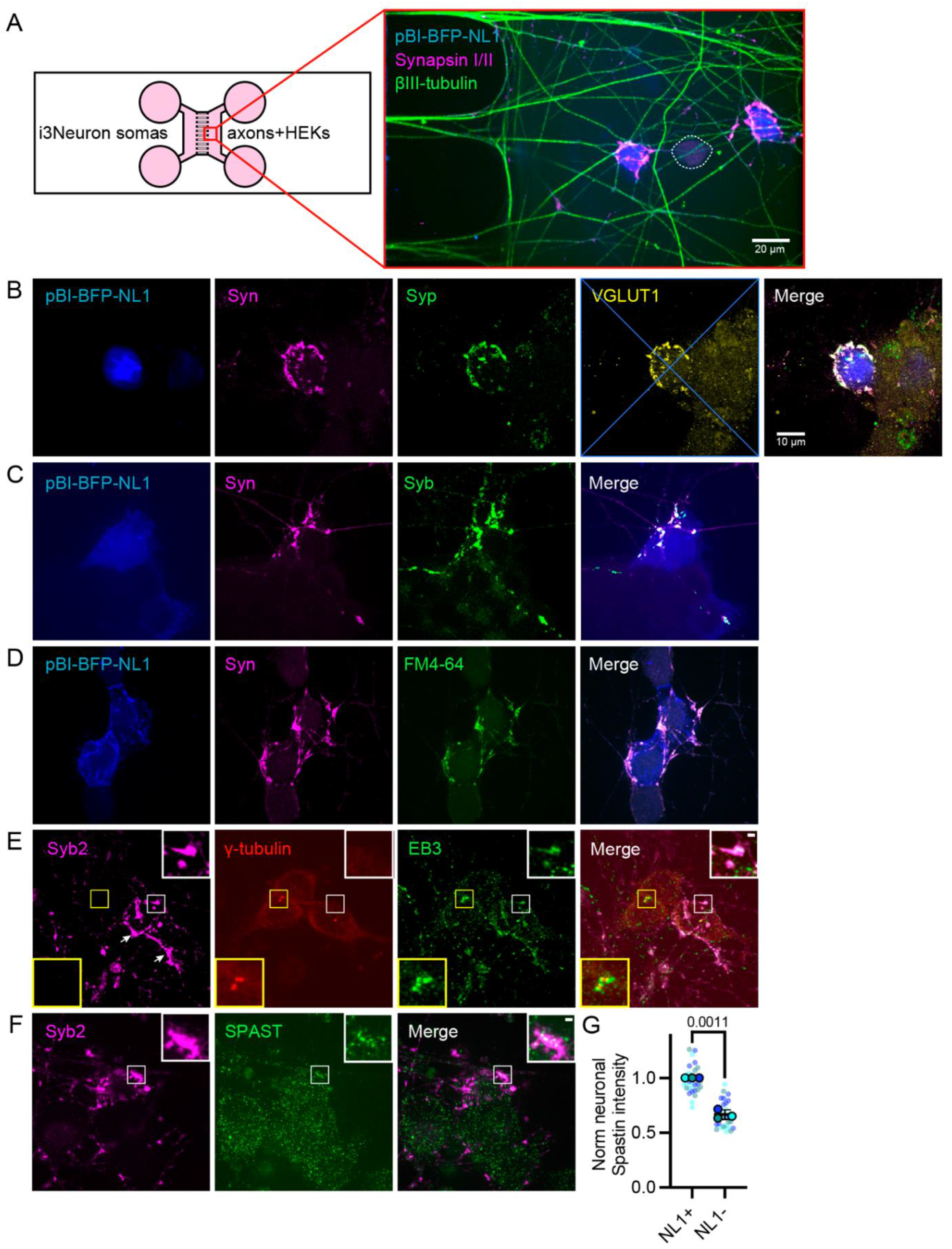
Human heterologous-culture assay generates presynapses containing cycling synaptic vesicles, +TIP EB3, and spastin. **(A)** Heterologous synapse model set-up with red inset showing βIII-tubulin+ axons forming Syn+ presynapses with NL1+ transfected HEK cells. Dotted white line emphasizes non-transfected HEK cell. Scale bar 20 µm. **(B-F)** Max intensity projections of i^3^Neuron axons crossing NL1+ HEK cells. Scale bar 10 µm. **(B-C)** Presynaptic markers Synaptophysin-1 (Syp; **B**), excitatory transmitter VGLUT1 **(B)**, and Synaptobrevin-2 (Syb; **C**) accumulate with Syn at heterologous synapses. **(D)** Syn+ heterologous synapses contain cycling synaptic vesicles as visualized by FM4-64 dye. **(E)** EB3, but not γ-tubulin, is enriched at Syb+ heterologous synaptic sites. Yellow box/inset reveals HEK γ-tubulin+ centrosomes nucleating EB3-decorated microtubules. White box/inset highlight presynaptic EB3. **(F)** Spastin is enriched at Syb+ presynapses. White box/inset highlight presynaptically enriched spastin at heterologous synapses. **(G)** Spastin intensity in heterologous synapses crossing NL1-expressing HEK cells (NL1+) and axonal regions crossing HEK cells not expressing NL1 (NL1-), normalized to spastin values in axons crossing NL1+ HEK cells. Error bars represent the standard deviation of the replicate means and provided *p*-values were determined by t-test comparing the means of three experimental replicates.

**Spastin regulates presynaptic component accumulation at heterologous presynapses.**

To determine whether spastin levels impact accumulation of presynaptic cargos, we employed the heterologous human synapse model to compare presynapse accumulation between CRISPRi spastin-targeting and non-targeting control i^3^Neurons (**Figure 8A-C**). Upon spastin depletion, we observed a decrease in the intensity of presynaptic components Syn (**Figure 8B**) and Syb (**Figure 8C**) 24 hours after NL1 introduction. Compellingly, spastin depletion altered accumulation of both presynaptic components despite Syn and Syb being trafficked via distinct mechanisms, with Syn undergoing slow axonal transport and Syb undergoing fast axonal transport by KIF1A [9, 34, 35]. These results reveal a specific reliance on spastin for priming new axonal regions for presynaptic zone formation upon contact with postsynaptic ligands.

**Figure 8.**
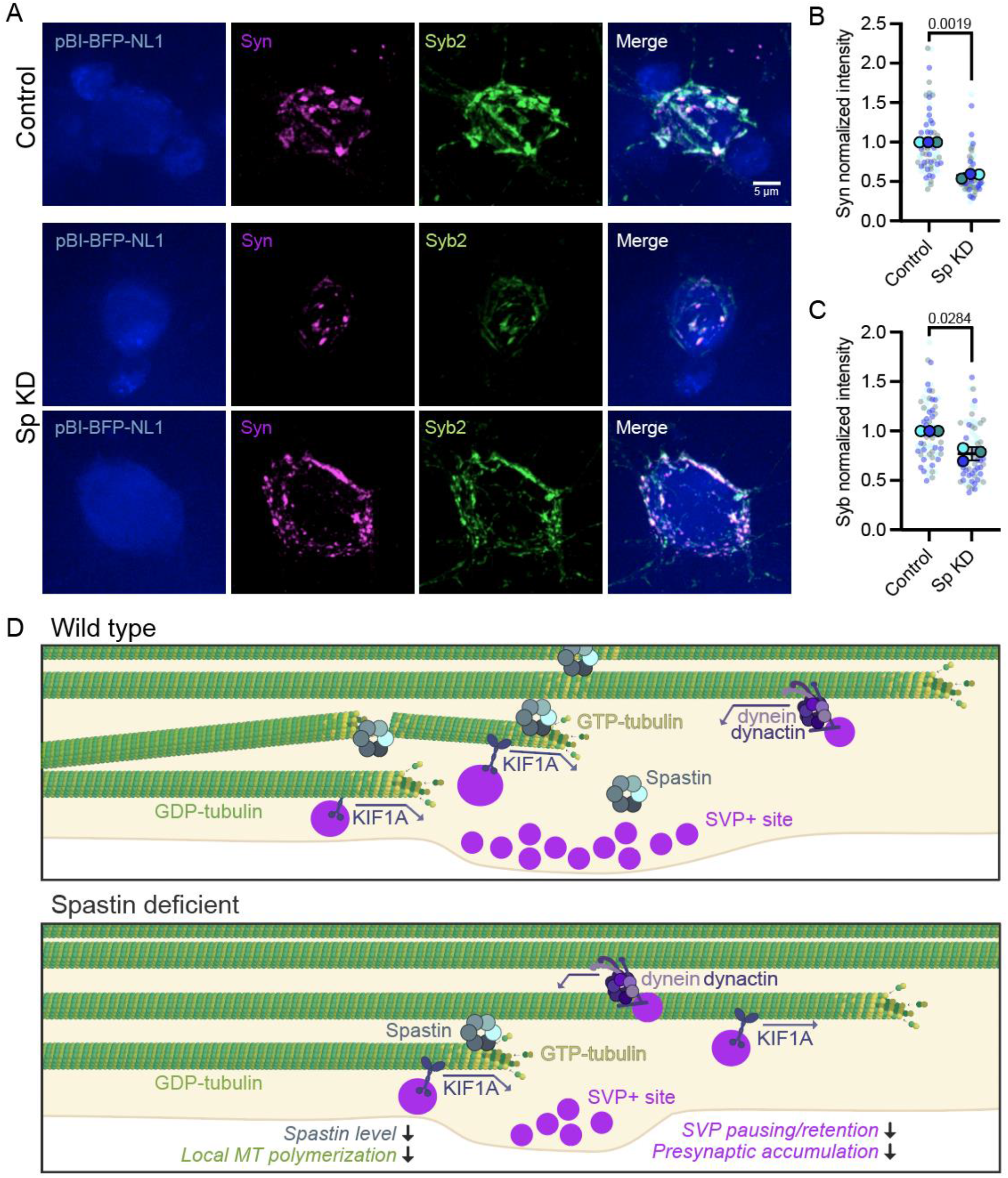
Spastin regulates presynaptic component accumulation. **(A)** Representative max intensity projections of CRISPRi non-targeting control (Control) and spastin CRISPRi-mediated knockdown (Sp KD) axons crossing pBI-BFP-NL1-expressing HEK cells (blue), stained for Syn (magenta) and Syb (green). Scale bar 5 µm. **(B-C)** Plots display Syn **(B)** and Syb **(C)** intensity values at heterologous synapses in Sp KD and Control axons normalized to Control. Error bars represent the standard deviation of the replicate means and provided *p*-values were determined by t-test comparing the means of three experimental replicates. **(D)** Model for spastin regulation of presynaptic cargo delivery. In wild-type neurons, presynaptic accumulations along the axon are enriched for spastin and exhibit increased localized microtubule polymerization. Spastin-mediated amplification of microtubule plus ends leads to increased anterograde SVP pausing/retention and overall presynaptic component accumulation. Upon spastin depletion, axons exhibit a decrease in localized microtubule growth events and reduced anterograde SVP pausing/retention. This results in fewer and less intense presynaptic accumulations in spastin knockdown neurons.

## DISCUSSION

We previously demonstrated that local microtubule dynamics sustain precise anterograde delivery of SVPs by KIF1A [7]. Here, we find that spastin locally modulates the microtubule landscape to promote polymerization at discrete sites, patterning the axon for SVP cargo pausing, retention, and overall presynaptic accumulation. Upon spastin reduction via CRISPRi, the preference for microtubule polymerization and SVP pauses/retention at SVP+ sites is diminished (**Figure 8D**), and presynaptic component accumulation at heterologous synapses is reduced. Our work supports a model where spastin locally enhances microtubule plus-end growth at specific sites along the axon to pattern synaptogenesis and guide presynaptic cargo distribution.

### Spastin-mediated microtubule polymerization influences presynaptic cargo pausing, retention, and accumulation along the axon

Recently, paradigm-shifting studies using reconstituted single-molecule assays demonstrated spastin’s potential to either depolymerize microtubules or increase microtubule mass [11, 12]. These *in vitro* experiments support cellular data revealing the paradoxical nature of spastin’s microtubule severing activity, with reduction of microtubule polymerization and/or mass observed upon either spastin overexpression or knockdown [13, 16–19]. Over the years, spastin’s severing activity has been implicated in numerous neuron-specific functions, including facilitating axon outgrowth [18, 20, 36], locally fragmenting microtubules in retreating motor axon branches [37], enhancing axon branch formation [36], regulating synaptic area in *Drosophila* neuromuscular junctions [38], and regulating CNS dendritic spine formation and maturation [21, 33]. These studies reveal the potential for spastin to undergo precise, circumstance-dependent regulation within neurons to influence a myriad of distinct functions.

We used human i^3^Neurons to uncover a novel role for neuronal spastin: locally enhancing microtubule plus-end amplification to direct presynaptic trafficking. Previous studies revealed the importance of localized microtubule polymerization at *en passant* presynaptic boutons in murine neurons, both for precise delivery of SVPs [7] as well as activity-induced exchange of synaptic vesicles [8]. In agreement with these findings, we found that presynaptic accumulations along human axons are hotspots of microtubule dynamics where microtubule polymerization preferentially initiates and terminates. Spastin is enriched at SVP+ sites and acts as important upstream regulator of localized microtubule growth. Our results further support a causal link between microtubule polymerization and SVP delivery, as spastin knockdown leads to both fewer microtubule comets and decreased anterograde pausing and retention of SVPs at SVP+ sites in i^3^Neuron axons.

Spastin depletion selectively inhibits anterograde SVP pausing and retention. Both anterograde-and retrograde-moving SVPs preferentially pause at SVP+ sites under normal conditions, but spastin KD only mildly affects retrograde trafficking behaviors. This suggests that retrograde SVP pausing/retention is mainly regulated independently of spastin-mediated microtubule modulation. However, spastin KD does decrease the percent of axons that experience preferential retrograde SVP+ pauses and leads to a trend toward fewer retrograde SVP+ retentions. This may be a consequence of dynein encountering fewer microtubule minus ends with decreased microtubule severing activity. While it is not yet possible to visualize individual microtubule filaments or naked minus ends within human axons, the observed spastin-mediated increase in microtubule plus ends (visualized as MACF43 comets) may correspond to an increase in microtubule minus ends when spastin severs microtubule filaments in two. If this is the case, however, one might expect a more dramatic retrograde response upon spastin depletion. In contrast to this model, spastin may act at SVP+ regions to generate microtubule lattice defects capable of promoting rescue (new bout of growth following a depolymerization event) as proposed by Vemu and colleagues [39]. In this scenario, spastin would promote localized microtubule polymerization without fully severing microtubules and generating new minus ends. This model is consistent with the observation of SVP pausing events at SVP+ regions not currently experiencing microtubule polymerization as well as concomitant with microtubule plus ends. Future live imaging and superresolution experiments may provide insight into which (if not both) of these models are in play at presynaptic sites along the axon.

Our human heterologous synapse assay brings together elements of previously employed mixed-culture synapse assays [40] and microfluidic axon isolation to dissect the role of spastin in presynapse formation. We discover that when i^3^Neuron axons are required to rapidly amass presynaptic components into heterologous presynapses, spastin KD inhibits the accumulation of two different presynaptic cargos, synapsin and synaptobrevin-2. Membrane-bound presynaptic components (including synaptobrevin-2 and synaptophysin), are packaged into SVPs and trafficked rapidly along the axon by KIF1A (200-400 mm/day; [9, 34, 35]). In contrast, synapsin undergoes slow axonal transport (2-8 mm/day; [34]). Reduced synaptobrevin-2 accumulation at spastin KD heterologous presynapses is consistent with our SVP live imaging experiments and in line with microtubule dynamics influencing KIF1A-mediated delivery [7]. However, the mechanism underlying the reduction of slow-moving synapsin upon spastin depletion is not yet clear. Both control and spastin KD axons are able to accumulate synapsin at heterologous presynapses within 24 hours of NL1 introduction, but spastin KD leads to dramatically lower endogenous synapsin intensity. This surprising result is consistent with the observation that synapsin moves anterogradely through stochastic, short-lived co-transport with synaptophysin-positive SVPs [41]. The reduction to synapsin accumulation upon spastin KD, therefore, could be a consequence of altered microtubule plus-end regulation of KIF1A. Future live imaging studies of human heterologous synapse formation may reveal how distinct motors, vesicle pools, and specific presynaptic components respond to presynaptic microtubule regulation.

Additional microtubule regulators may also influence presynaptic microtubule dynamics. The microtubule nucleator γ-tubulin, for example, has been implicated in activity-dependent microtubule nucleation at presynaptic boutons in primary rat neurons [8]. However, γ-tubulin is not observed at human presynapses in our heterologous culture model. We also find that human i^3^Neurons contain fewer axonal microtubule growth events compared to rat primary cortical neurons. i^3^Neurons at DIV21 exhibit one third the average number of axonal comet events compared to DIV11 rat primary cortical neurons as determined by tracking GFP-MACF43 [42]. These divergences may reflect species-specific regulation or synapse maturation state/activity differences between the culture models. Spastin severing activity is also regulated by the microtubule post-translational modification (PTM) polyglutamylation [17, 38, 43] which is the addition of multiple glutamates to C-terminal tubulin tails [44]. *In vitro* reconstituted single molecule assays reveal that polyglutamylation tunes spastin severing as a function of glutamates per tubulin but becomes inhibitory past a threshold [43]. Neuronal microtubules are highly polyglutamylated [45], but it is challenging experimentally to determine whether/how polyglutamylation level varies across the neuron. Spastin levels have previously been linked to amount of neuronal tubulin polyglutamylation [21, 37, 38], suggesting an interesting feedback between the two molecular players. Advancements to superresolution microscopy techniques may shed light on the physiological relationship (if any) between spastin and polyglutamylation at synaptic sites.

### Spastin mutations associated with synaptic dysfunction are clustered in the functional AAA domain essential for severing activity

The importance of spastin in neural connectivity is bolstered by recent reports of human patients with mutations to spastin exhibiting synapse-related deficiencies including memory impairment, intellectual disability, seizures, autism spectrum disorder, and severe depression [22]. Spastin, encoded by the gene *SPG4*, was originally discovered by mapping mutations in patients presenting with autosomal dominant hereditary spastic paraplegia (HSP) [46]. HSP is a neurodegenerative disorder characterized by lower limb spasticity and weakness caused by progressive distal degeneration of the corticospinal tract. The etiology of spastin-associated HSP is still debated but has been proposed to be a combination of haploinsufficiency and toxic gain-of-function mechanisms [47]. Pathology is complicated by the presence of multiple spastin isoforms with distinct cellular regulation and numerous mutation types including heterozygous missense, nonsense, truncating, and frameshift [48]. The M1 isoform, derived from translation starting at methionine 1, is mainly expressed in the adult spinal cord [27], contains an N-terminal hairloop domain that targets it to endoplasmic reticulum (ER) membrane and lipid droplets [49], and is thought to contribute little to microtubule severing [47]. The M87 isoform, derived from translation starting at methionine 87, is the primary isoform expressed in the developing nervous system [27], lacks the N-terminal hairloop domain, and is responsible for microtubule severing throughout the neuron. M1 is expressed in the right place (corticospinal tract) at the right time (adulthood), for gain-of-function mutants to contribute strongly to HSP neurodegeneration [47]. In contrast, spastin-associated synaptic deficiencies are often linked to mutations to the AAA functional domain [22, 50]. The vast majority of spastin-associated psychiatric manifestations result from truncating mutations to the AAA domain [22], and intellectual disability [50] and memory impairment [22] are more prevalent in cases of AAA missense mutations. In combination with previous reports of spastin acting at the postsynapse [21, 33] and our new findings revealing a role for spastin presynaptically, these phenotype-genotype analyses highlight the importance of AAA-ATPase-mediated microtubule severing in synapse development and function.

### Protosynapse formation and maintenance may represent an upstream step in neuron synaptogenesis

Characterization of i^3^Neuron synapses yielded the unexpected result that most presynaptic puncta within human axons at DIV21 in monoculture are not apposed by postsynaptic densities. In fact, over 80% of presynaptic puncta are ‘protosynapses’ lacking a postsynaptic partner. By extending i^3^Neuron maturation to DIV42 by co-culture with primary rat astrocytes, the percentage of presynaptic puncta that are apposed by postsynaptic partners increases dramatically and a nearly five-fold increase in synapse density is observed. Both *bona fide* presynapses and protosynapses are enriched for endogenous microtubule +TIP EB3 and spastin, consistent with our live imaging results at stable SVP+ sites. Of note, both *bona fide* presynapses and protosynapses are capable of cycling SVs as shown by uptake of FM4-64 dye. This aligns with previous reports of neurons establishing functional presynaptic zones independent of postsynaptic contacts both in culture [5] and *in vivo* [4], and the rapid rate at which SVPs and active zone proteins coalesce into functional synaptic puncta behind the growth cone in *C. elegans* PDE axons [51]. As dopaminergic PDE neurons do not require strict apposition with postsynaptic densities [51], this result reveals an *in vivo* mechanism of presynapse establishment in newly constructed axons independent of specific postsynaptic interaction. In human neurons, axonal presynaptic component clustering may therefore represent an upstream step in synaptogenesis. In this pre-patterning model, presynaptic machinery would be interspersed along the axon, ready to be stabilized and matured upon axodendritic contact. This model is further supported by the observation that dendritic filopodia preferentially form stable contacts with SVP+ axonal sites [6]. It is also consistent with the observed transition from protosynapse *to bona fide* presynapse upon i^3^Neuron maturation. Taken together, our pre-patterning model advocates for the structural landscape of the axon, the microtubules themselves, priming specific regions of the axon for synaptogenesis.

Interestingly, the density of stable SVP+ sites (determined using live imaging data) is similar to the density of axonal Syn+ protosynapses (determined in microfluidically isolated axons) and heterologous presynapses (determined within single axons crossing NL1+ HEK cells), with all approximately 1 puncta/10 µm of axon. We observe that i^3^Neuron axons typically host stretches of protosynapses that are generally smaller and further apart than heterologous presynapses, which may reflect a mobilization and coalescing of puncta upon axonal contact with postsynaptic ligand. It is not clear why the heterologous synapse does not occupy the entire stretch of axon experiencing NL1, but the observed patterning suggests a cellular mechanism in place to limit presynapse size and set spacing. Future experiments marrying the heterologous synapse assay with live-imaging approaches may shed light on whether/how protosynapses are mobilized or stabilized upon postsynaptic contact.

If protosynapse establishment occurs prior to initial postsynaptic contact, what dictates protosynapse formation? With spastin enriched at the distal axon and the knowledge that presynaptic puncta can form rapidly behind an extending growth cone [51], it is tempting to speculate that the spastin-mediated SVP targeting mechanism reported here may also influence protosynapse formation in the nascent axon. Live imaging experiments in advancing growth cones of control and spastin-depleted axons may shed light on whether this supposition is correct. Probing this question *in vivo*, where neurons experience numerous cell types and external cues, will also be essential in understanding physiological regulation of axonal presynaptic patterning.

## CONCLUSIONS

Our study provides new insight into the cellular mechanisms regulating presynaptic component accumulation along the axon, placing the microtubule severing enzyme spastin upstream of local presynaptic microtubule polymerization. Using human i^3^Neurons, we find that specific axonal regions housing stable accumulations of presynaptic components are hotspots both for microtubule polymerization events and SVP pausing in the anterograde and retrograde direction. Disruption of neuronal spastin, either by depletion via CRISPRi or by transient overexpression, interrupts the localized enrichment of microtubule amplification and diminishes SVP accumulation. These findings support a model where spastin locally enhances microtubule growth to guide presynaptic cargo delivery and pattern the axon for synaptogenesis.

## ACKNOWLEDGEMENTS

We thank Mariko Tokito for assistance with cloning the expression plasmid constructs. We are grateful to Ruud Toonen from Vrije Universiteit Amsterdam for sharing protocols and providing guidance on co-culturing iNeurons on primary rat astrocytes, as well as Caela Long and Judy Grinspan from the University of Pennsylvania for generously assisting with isolation of primary rat astroglial cells. We thank Carris Borland, Bishal Basak, and Dan Dou for insights and discussions. The authors gratefully acknowledge support from NIH awards to J.A. (NINDS F32NS117672) and E.L.F.H. (NIGMS R35 GM126950).

## AUTHOR CONTRIBUTIONS

J.A. and E.L.F.H. conceived the project and designed experiments. J.A. performed the experiments, analyzed the data, and wrote the manuscript with contributions from E.L.F.H.

## STAR METHODS

### RESOURCE AVAILABILITY

#### Lead contact

Further information and requests for resources and reagents should be directed to and will be fulfilled by the lead contact, Erika Holzbaur.

#### Materials availability

All cell lines and plasmids generated in this study are available upon request.

#### Data and code availability

Excel datasets for all graphs will be uploaded to Xenodo. No new code was generated for this study.

### EXPERIMENTAL MODEL AND SUBJECT DETAILS

#### Human induced pluripotent stem cells (iPSCs)

Human iPSCs (male, WTC11 background [52]) were cultured in 10 ml Essential 8 (E8) Medium (Gibco™/Thermo Fisher Scientific; A1517001) on standard 10 cm sterile tissue culture plates treated with Matrigel Growth Factor Reduced, LDEV-Free Basement Membrane Matrix (Corning®; 356231) diluted 1:50 in DMEM/F-12 (Gibco™/Thermo Fisher Scientific; 11320-033). Cells were incubated at 37C and 5% CO_2_ in a water-jacketed cell culture incubator and were passaged upon reaching 60%–80% confluency. The following steps were performed for each passage: cells were washed twice with PBS, pH 7.4 (Gibco™/Thermo Fisher Scientific; 10010-23), incubated with 3 ml Accutase® solution (Sigma-Aldrich; A6964) in 37C incubator for 5-6 min, checked for cell dissociation, equal volume E8 media supplemented with 10nM Y-27632 dihydrochloride ROCK inhibitor (Tocris; 125410) was added, cells were collected in 15 ml conical, centrifuged 200 g for 5 min, resuspended in E8 media supplemented with 10nM ROCK inhibitor, counted using Countess™ Automated Cell Counter, and finally plated at 6-8×10^5^ cells per plate into new, Matrigel-treated 10 cm tissue culture plates. E8 media was changed daily.

#### Human iPSC-derived i^3^Neurons

Human iPSC-derived i^3^Neurons were differentiated from human WTC11 iPSCs engineered to express NGN2 under a doxycycline-inducible system in the AAVS1 safe harbor locus [24, 30] and pC13N-dCas9-BFP-KRAB targeted to the human CLYBL intragenic safe harbor locus [29]. Differentiation of iPSCs into i^3^Neurons was performed using an established protocol [30] involving the following steps: release iPSCs using Accutase® solution and centrifuge as above, resuspend pelleted cells in Induction Media consisting of DMEM/F-12 (Gibco™/Thermo Fisher Scientific; 11330-032) supplemented with 1X MEM Non-Essential Amino Acids (NEAA) (Gibco™/Thermo Fisher Scientific; 11140-050), 1X N2 Supplement (Gibco™/ Thermo Fisher Scientific; 17502-048), 1X GlutaMAX™ Supplement (Gibco™/Thermo Fisher Scientific; 35050-061), 10 nM ROCK inhibitor, and 2 mg/mL doxycycline (Sigma; D9891) to induce expression of mNGN2. iPSCs were counted using Countess™ Automated Cell Counter and plated at 5-7×10^5^ cells per Matrigel-coated 15 cm dish in 20 mL of Induction Media. After 24 and 48 hours, media was changed to 20 ml Induction Media without ROCK inhibitor. 72 hours after plating, pre-differentiated cells were cryopreserved in liquid N_2_ in Cryopreservation Media consisting of BrainPhys™ Neuronal Medium (STEMCELL Technolgies; 05790) supplemented with 2% B27 Supplement (Gibco™/Thermo Fisher Scientific; 17504-044), 10 ng/mL BDNF (PeproTech; 450-02), 10 ng/mL NT-3 (PeproTech; 450-03), 1 mg/mL Mouse Laminin (Thermo Fisher Scientific; 23017-015), 10% Dimethylsulfoxide (DMSO;), and 20% Fetal Bovine Serum (FBS; HyClone; SH30071.03). The published i^3^Neuron differentiation protocol can be found on Protocols.io (https://doi.org/10.17504/protocols.io.261ge348yl47/v1).

#### Generation of CRISPRi spastin-targeting or non-targeting control iPSCs

Spastin-specific sgRNA guide GACCGACGGGAACCAAGCGA or non-targeting sgRNA guide GTGCCAGCTTGTGGTGTCGT was cloned into pCRISPRia-vs2 ([53]; Addgene; Plasmid 84832) using previously described methods [54]. Briefly, the guides were flanked with BstXI and BlpI cutsites and ligated into BstXI/BlpI-digested pCRISPRia-vs2 using NEB Quick Ligase (New England Biolabs; M2200S). Guide incorporation was verified by sequencing using the Forward and Reverse oligonucleotides ggcttggatttctataacttcg and ctactgcacttatatacggttc, respectively. The sgRNA guides were then packaged into lentivirus and transduced into iPSCs based on a previously published protocol [29] using the following steps: 15 cm culture dishes were seeded with 8×10^6^ HEK293T cells in 20 ml Dulbecco’s Modified Eagle’s Medium (DMEM; Corning; 10-017-CM), which was supplemented with 10% FBS (HyClone; SH30071.03). After 24 hours, the cells were transfected with a 1:1:1 mix of 2.5 µg sgRNA plasmid, psPAX2 (HIV pol+gag), pCMV-Vsv-g with Lipofectamine 2000 Transfection Reagent (Invitrogen; 11668027). Eight hours following addition of transfection solution, media was replaced with 20 ml Growth Medium supplemented with 1:500 dilution ViralBoost (Alstem; VB100). Two days following transfection, cell media was collected, filtered, and centrifuged with Lentivirus Precipitation Solution (Alstem; VC100) to isolate viral pellet. The virus-containing pellet was resuspended in 10 mL E8 media with ROCK inhibitor, aliquoted, and stored at −80C. For viral transduction, 8×10^5^ iPSCs were seeded into T25 cell culture flasks with 2 ml thawed E8 media containing virus and ROCK inhibitor and incubated for six hours before adding an additional 3 ml E8 supplemented with ROCK inhibitor. Two days following virus introduction, iPSCs were passaged (1.5×10^6^ cells per 10 cm plate) and cultured in 10 ml E8 supplemented with ROCK inhibitor and 0.8 µg/mL puromycin (Takara; 631305). After four days of replacing media with E8 supplemented with 1 µg/mL puromycin, cells were collected and cryopreserved in liquid N_2_.

#### Culture and transfection of iPSC-derived neurons in monoculture

Cryopreserved, pre-differentiated neurons (i^3^Neurons or CRISPRi i^3^Neurons) were thawed and plated on 35-mm glass-bottom imaging dishes (MatTek; P35G-1.5-20-C) coated in 100 µg/mL poly-L-ornithine (Sigma-Aldrich; P3655) at densities of 3-5×10^5^ cells per dish. At least two different neural induction batches were used in three or four independent experimental cultures for each condition. iPSC-derived neurons were cultured in i^3^Neuron Maintenance Media consisting of BrainPhys™ Neuronal Medium (STEMCELL Technolgies; 05790) supplemented with 2% B27 Supplement (Gibco™/Thermo Fisher Scientific; 17504-044), 10 ng/mL BDNF (PeproTech; 450-02), 10 ng/mL NT-3 (PeproTech; 450-03), and 1 mg/mL Mouse Laminin (Thermo Fisher Scientific; 23017-015). Once a week, 40% of the media was replaced with fresh, pre-equilibrated Neuron Maintenance Media. Live imaging and immunocytochemistry experiments were performed 21 days after thawing pre-differentiated iPSC-derived neurons (DIV21), unless otherwise noted. Most experiments were performed at DIV21 due to the tendency of i^3^Neurons to detach from the plate with prolonged culture. For live imaging, i^3^Neurons were transfected with Lipofectamine Stem Transfection Reagent (Thermo Fisher Scientific; STEM00001) and 0.75-2 μg total plasmid DNA. Published protocol for culture and transfection of iPSC-derived neurons for live-imaging of axonal cargos can be found on Protocols.io (https://doi.org/10.17504/protocols.io.x54v9dj4zg3e/v1).

#### Rat primary astrocyte isolation and culture

All experiments were performed in accordance with the guidelines set forth by The Children’s Hospital of Philadelphia and The University of Pennsylvania Institutional Animal Care and Use Committees. Primary rat astrocytes were isolated from brains of Sprague Dawley rats (Charles River Laboratories, Wilmington, MA RRID: RGD_737891) at postnatal day 1 and plated on poly-D-lysine-coated T75 flasks and cultured in Neurobasal medium (Gibco™/Thermo Fisher Scientific; 21103049) with 2% B27 supplement at 37°C with 5% CO_2_ as previously described [55]. After 24 hours, growth medium was changed to Neurobasal medium with B27 and 10 ng/ml bFGF, 2 ng/ml PDGF-AA, and 1 ng/ml neurotrophin-3. Once cells reached confluence (around DIV7), astroglia cells were separated from oligodendroglial cells using the “shake-off” method [56], and remaining astroglial were split into new T75 flasks using Trypsin-EDTA (Gibco™/Thermo Fisher Scientific; 25300-054) and cultured in astroglia growth media consisting of DMEM (Corning; 10-017-CM) with 10% FBS (HyClone; SH30071.03), 1% NEAA, and 1% Pen/Strep. Media was changed weekly until cells reached confluence (approximately DIV14). Primary rat astrocytes were then frozen in cryopreservation media (astroglia growth media with 10% DMSO).

#### Co-culture of iPSC-derived neurons with primary rat astrocytes

Primary rat astrocytes were thawed into uncoated T75 flasks and cultured in astrocyte growth media at 37°C with 5% CO_2_. Once confluent (around 7 days post thaw), cells were passaged into 35-mm glass-bottom imaging dishes using Trypsin-EDTA at a density of 50,000 cells per dish.

Once confluent (approximately 4-7 days post passage), 50,000 cryopreserved, pre-differentiated i^3^Neurons were thawed and plated on the 35-mm glass-bottom imaging dishes pre-seeded with primary rat astrocytes and cultured in pre-equilibrated Neuron Maintenance Media. Once a week, 40% of the media was replaced with fresh, pre-equilibrated Neuron Maintenance Media. Immunocytochemistry experiments were performed 21 days or 42 days after thawing pre-differentiated iPSC-derived neurons (DIV21 or DIV42, respectively).

#### Human embryonic kidney cells

Human embryonic kidney cells (HEK293T, referred to as HEK cells) were authenticated by STR profiling. HEKs cells were cultured in DMEM (Corning; 10-017-CM), which was supplemented with 10% FBS on standard 10 cm sterile tissue culture plates at 37C in 5% CO_2_ in a water-jacketed cell culture incubator. Cells were passaged using trypsin upon reaching 60%–90% confluency.

## METHOD DETAILS

### Molecular Cloning

Expression constructs used in this study include pCI-mScarlet (Addgene #85042; pC1-mScarlet-Syp [7]), pCIG2-GFP-MACF43 [42], pEGFP-C3-Syb (Addgene #42308; used to make pmScarlet-Syb), pCIG2-mScarlet-Syb-IRES-GFP-MACF43 (generated for this study by inserting mScarlet-Syb into pGIC2-GFP-MACF43 using sticky-end cloning at XbaI/PspXI), pCMV-Tag/WT M87 (Addgene #89322), pCIG2-SNAP-M87 (generated for this study from pCIG2-GFP-MACF43 by replacing GFP with SNAP-tag from pSNAPf vector (New England Biolabs) and MACF43 with M87 from pCMV-Tag/WT M87), pBI-BFP (made by replacing AcGFP1 in pBI-CMV2 Vector (Clontech) with BFP from pTagBFP-N Vector (Evrogen) at EcoRI/XbaI), pCAG-NL1 (Addgene #15260), and pBI-NL1-BFP (generated for this study by inserting NL1 into pBI-BFP using sticky-end cloning at NheI/EcoRV). All constructs were verified by DNA sequencing.

### Live imaging of i^3^Neurons

i^3^Neurons plated on 35-mm glass-bottom imaging dishes were imaged in an environmental chamber at 37C in i^3^Neuron Imaging Media consisting of low fluorescence Hibernate A Medium (Brain Bits®; HA100) supplemented with 2% B27, 10 ng/mL BDNF, 10 ng/mL NT-3, and 1 mg/mL Mouse Laminin. Time-lapse imaging data presented in Figure 1 were acquired on a PerkinElmer UltraView Vox Spinning Disk Confocal system with a Nikon Eclipse Ti inverted microscope, using a Plan Apochromat 60× 1.40 NA oil immersion objective and a Hamamatsu EMCCD C9100 −50 camera controlled by Volocity software. Following a scheduled microscope upgrade, all other imaging data was acquired using a Hamamatsu ORCA-Fusion C14440-20UP camera controlled by VisiView software. Imaged axonal regions were selected based on morphological parameters: long processes of uniform diameter without noticeable swellings at least ∼200 μm away from the cell body. For spastin SNAP-M87-expressing neurons, cells were incubated with SNAP-tag ligand SNAP-Cell 647-SiR (New England BioLabs; S9102S) at a final concentration of 100 nM for 15 minutes followed by 30-45 minute washout in i^3^Neuron Maintenance Media before imaging in i^3^Neuron Imaging Media. For SVP motility experiments and spastin M87-overexpression neurons, a one-channel (mScarlet-Syp/Syb for SVP motility experiments) or three-channel (MACF43-GFP, mScarlet-Syb, and SNAP-M87 for spastin overexpression) z-stack was acquired prior to live imaging to help determine position of static Syp/Syb and/or spastin puncta. To aid the visualization and tracking of mobile SVPs, the whole image field was photobleached with the 561 nm laser at full power for 10 seconds before time-lapse imaging began. Time-lapse images of SVPs (mScarlet-Synaptophysin or mScarlet-Synaptobrevin-2) were then collected at a frame rate of 200 ms per frame for 4 minutes. For time-lapse imaging of microtubule comets, two-channel acquisition of MACF43-GFP and mScarlet-Syp/Syb was performed at a frame rate of 1 second per frame for 5 minutes.

### Heterologous synapse assay

XonaChip microfluidic chambers with 450 µm barriers (Xona Microfluidics; XC450) were coated with Xona Pre-Coat Solution for 13 min at room temperature, washed twice with PBS, coated with 20 µg/mL poly-L-ornithine at 4C overnight, washed with ddH_2_O, and coated with 20 µg/mL Mouse Laminin at 4C overnight (following manufacturer instruction to ensure proper fluid flow at each step). After washing with PBS, pre-equilibrated i^3^Neuron Maintenance Media (described above) was added to each chamber and the device was further pre-equilibrated at 37C in 5% CO_2_ for >30 minutes. Cryopreserved, pre-differentiated neurons (i^3^Neurons or CRISPRi i^3^Neurons) were thawed and 80,000 cells were seeded into one chamber of the coated, pre-equilibrated device following manufacturer instructions. After 24 hours, media was exchanged with fresh, pre-equilibrated i^3^Neuron Maintenance Media. Pre-equilibrated i^3^Neuron Maintenance Media was added to each well twice per week. At DIV13, HEK cells transfected 24 hours prior with 6 µL:1 µg mix of FUGENE and pBI-NL1-BFP (bicistronic vector expressing untagged NL1 and cytosolic BFP) were added to the axonal chamber of the XonaChip. XonaChips containing DIV14 i^3^Neurons and HEK cells were fixed for immunocytochemistry 24 hours after HEK cell addition.

### FM4-64 synaptic cycling assay

For FM-dye labeling of active synaptic vesicle cycling, cells were exposed to 10 μM SynaptoRed™ C2 (Equivalent to FM®4-64; Biotium; 70027) in Stimulation Buffer consisting of 31.5 mM NaCl, 90 mM KCl, 5 mM Hepes, 1 mM MgCl_2_, 2 mM CaCl_2_, 30 mM glucose, and 50 μM D-AP5 (Thomas Scientific; 14539-5) for 2 min. Cells were then incubated with 10 μM SynaptoRed™ C2 in Wash Buffer consisting of 50 µM D-AP5 and 10 µM CNQX in Hibernate A for 5 min. Cells were washed with Wash Buffer alone, Wash Buffer with 1 mM ADVASEP-7 (Biotium; 70029), and finally Wash Buffer alone. Cells were then fixed in 4% paraformaldehyde/4% sucrose in PBS for 10 min at room temperature and immunostained for synaptic markers (details below).

### Immunocytochemistry of i^3^Neurons

For immunocytochemistry of synapses or heterologous synapses, DIV21 Human i^3^Neurons plated in 35-mm glass-bottom imaging dishes or XonaChips were fixed in 4% paraformaldehyde/4% sucrose in PBS for 10 min at room temperature. For XonaChips, solutions were added following manufacturer instruction to ensure proper fluid flow at each step. Following three PBS washes, cells were permeabilized/blocked in Blocking Solution (1% bovine serum albumin (BSA), 5% goat serum in PBS) with 0.1% Triton X-100 for 1 hour. Neurons were then incubated with primary antibodies for proteins of interest diluted in Blocking Solution at 4C overnight. Primary antibodies used with 4% paraformaldehyde/4% sucrose fixation include Synapsin I/II (anti-guinea pig; Synaptic Systems: 106 004; 1:1000 dilution), PSD-95 (anti-rabbit; Synaptic Systems: 124-008-SY; 1:500 dilution), MAP2 (anti-mouse; EMD Millipore; MAB3418; 1:200 dilution), VGLUT1 (anti-goat; Synaptic Systems; 135 307; 1:500 dilution), Synaptophysin(anti-mouse; Millipore Sigma; S5768; 1:200 dilution), and Synaptobrevin-2 (anti-rabbit; Cell Signaling; 13508; 1:250 dilution). Following three washes with PBS, neurons were incubated protected from light with species-matched secondary antibodies (1:500 dilution) diluted in blocking solution for 1 hour at room temperature. After three washes with PBS, coverslips were mounted in ProLong Gold Antifade Mountant (Thermo Fisher;). Images were acquired as z-stacks with 200 nm step-size using the Hamamatsu ORCA-Fusion C14440-20UP camera controlled by VisiView software as described above.

For immunocytochemistry of synaptic spastin, EB3, and γ-tubulin in DIV21 Human i^3^Neurons plated in 35-mm glass-bottom imaging dishes or XonaChips, cells were washed once with warm PHEM buffer (60 mM PIPES, 25 mM HEPES, 10 mM EGTA, 2 mM Mg2Cl) and then fixed with ice-cold methanol for 5 min at −20C. After fixation, cells were permeabilized with 0.1% Tween 20 in PBS for 3 minutes and blocked in Blocking Solution (1% BSA, 5% goat serum in PBS) with 0.1% Triton X-100 for 30 minutes. Neurons were incubated with primary antibodies against spastin 3G11/1 (anti-mouse; Santa Cruz Biotechnology; sc-53443 1:250 dilution), EB3 (anti-rat; Abcam; ab53360; 1:100 dilution), and/or γ-tubulin (anti-mouse; Millipore Sigma; T6557; 1:1000 dilution) with Synapsin I/II, PSD-95, and MAP2 (MAP2 not used with spastin immunostaining due to species overlap) or Synaptobrevin-2 diluted in Blocking Solution at 4C overnight. Following three washes with PBS, neurons were incubated with secondary antibodies diluted in blocking solution for 1 hour at room temperature and mounted and imaged as above.

### Immunoblotting

i^3^Neuron samples were lysed in RIPA buffer (50 mM Tris-HCl, 150 mM NaCl, 0.1% Triton X-100, 0.5% deoxycholate, 0.1% SDS, 2× Halt Protease and Phosphatase inhibitor), centrifuged for 15 min at 18,000 to clear cellular debris, and protein concentration of the supernatant was determined by Pierce™ BCA Protein Assay Kit (Thermo Fisher Scientific; 23225). Spastin protein was resolved on 8% SDS-PAGE gels transferred to Immobilon®-FL PVDF membranes (Millipore; 05317) using a wet blot transfer system (BioRad). Membranes were then stained for total protein using LI-COR Revert 700 Total Protein Stain and imaged using an Odyssey CLx Infrared Imaging System (LI-COR). Following imaging of total protein stain, membranes were de-stained and blocked for 5 minutes with EveryBlot Blocking Buffer (BIO-RAD; 12010020). Membranes were incubated with spastin primary antibody (Sp 3G11/1; Santa Cruz Biotechnology; sc-53443) diluted 1:200 in EveryBlot at 4C overnight. Membranes were washed three times in TBS (50 mM Tris-HCl [pH 7.4], 274 mM NaCl, 9 mM KCl) with 0.1% Tween-2 and incubated with mouse secondary antibodies diluted 1:20,000 in EveryBlot with 0.01% SDS for 1 hr at room temperature. Following three washes with TBS with 0.1% Tween-20 and one wash with TBS, membranes were imaged on the Odyssey CLx Infrared Imaging System. Western blots were analyzed with Image Studio Software (LI-COR) comparing level of spastin protein standardized to total protein in Control vs. Spastin-targeting CRISPRi i3Neurons. Published protocol can be found on Protocols.io (https://doi.org/10.17504/protocols.io.5jyl8j5zrg2w/v1). N=3 experimental replicates from 3 separate i^3^Neuron inductions.

## QUANTIFICATION AND STATISTICAL ANALYSIS

### Kymograph generation and tracing

Kymographs of live-imaging series were generated using the KymographClear plugin for Fiji [57] following published instructions [58] with line width set to 3 pixels. Axonal regions of ∼100 µm with active SVP trafficking and/or microtubule comets were selected for quantification. Within the kymograph, individual tracks were traced manually in Fiji and the coordinates of each inflection point were input into an excel sheet. Coordinates were then converted from pixels to µm (x-coordinate) and seconds (y-coordinate) for further analysis. The distance (x) and duration (y) values were then used to calculate SVP movement or microtubule comet parameters. SVP+ sites and lengths were determined based on the location of bright immobile clusters of either mScarlet-Syp or mScarlet-Syb along the axon.

### SVP movement analysis

Motile SVPs were scored as anterograde (net displacement >10 µm in the anterograde direction) or retrograde (net displacement >10 µm in the retrograde direction). SVP flux of vesicles per minute was calculated as the number of anterograde or retrograde SVPs entering the kymograph normalized to imaging time. The coordinates of SVP tracks were used to calculate instantaneous velocity, pause frequency, and pause duration. A pause was defined as a single or consecutive instantaneous velocity value of <0.083 mm/s lasting at least 1 second in duration. A retention event was defined as an SVP that moved into the kymograph, stopped, and remained stationary for the remainder of the acquisition (SVP had to be stationary for at least the last 30 seconds of the kymograph to be counted as a retention). SVP+ regions were determined by comparing Syp/Syb puncta location in initial z-stack and the position/intensity of stable (vertical) signal of Syp/Syb in the kymograph.

SVP pause frequency (pauses/SVP/µm/min) in SVP+ regions was determined by normalizing the number of SVP pauses in the anterograde or retrograde direction occurring within SVP+ regions by the total number of anterograde or retrograde SVPs, stable SVP coverage of the axon, and imaging time. SVP pause frequency in SVP-regions was determined by normalizing the number of SVP pauses in the anterograde or retrograde direction occurring outside of SVP+ regions by the total number of anterograde or retrograde SVPs, non-synaptic coverage of the axon, and imaging time. SVP+/SVP-ratio for each analyzed axon was determined by dividing the SVP+ pause frequency by the SVP-pause frequency, and the percent of axons that fell into the ratio bin of 0-1, 1-5, and 5+ were plotted. N=4 experimental replicates, n≥27 axons.

### Microtubule comet analysis

The coordinates of microtubule comet traces were used to determine average polymerization rate (comet distance/duration), microtubule polymer added (total comet distance/kymograph distance/imaging time), microtubule comet density (number of comets/10 µm/imaging time), polymerization distance, and polymerization time. SVP+ regions were determined by the position/intensity of stable (vertical) signal of Syp/Syb signal in mScarlet-Syp/Syb kymographs. SVP+ and SVP-microtubule comets, initiation events, and termination events were determined manually by counting the number of events that occur within SVP+ sites (SVP+) or elsewhere in the axon (SVP-). Initiation or termination frequencies (events/10 µm/minute) in SVP+ regions were determined by normalizing the number of events occurring within SVP+ regions by SVP+ coverage of the axon and imaging time. Initiation or termination frequencies (events/10 µm/minute) in SVP-regions were determined by normalizing the number of events occurring outside of SVP+ regions by distance of axon not stably inhabited by SVPs and imaging time. SVP+/SVP-ratio for each analyzed axon was determined by dividing the SVP+ initiation or termination frequency by the SVP-initiation or termination frequency, and the percent of axons that fell into the ratio bin of 0-1, 1-5, and 5+ were plotted. N=4 experimental replicates, n≥27 axons.

### Kymograph SVP+ site intensity analysis

Mean SVP+ site intensity was determined using Fiji by measuring mScarlet-Syb intensity for size matched region of interest (ROI) encompassing stable SVP+ regions. SVP+ ROI intensity was adjusted for background signal by subtracting background fluorescence intensity measurement of the same size. Adjusted SVP+ intensity values were normalized to replicate control average, and normalized cell averages and experiment averages were plotted. N=4 experimental replicates, n≥27 axons.

### SynapseJ analysis

Z-stack images of i^3^Neurons at DIV14, DIV21 in monoculture or DIV21, DIV42 co-cultured with primary rat astrocytes were immunostained for Synapsin I/II, PDS-95, and MAP2 and analyzed for synapse detection using SynapseJ (v.1) following published instructions [31]. Detection threshold was set to one-half the mean puncta intensity for pre- and postsynaptic channel (determined using 3D objects counter) and default noise, puncta size, and Find Maxima were applied. Following SynapseJ automated synapse identification, SynapseJ merge.tif files were opened and Fiji’s 3D Object Counter was used to identify number of apposed pre- and postsynaptic puncta from the 3D masks generated in SynapseJ. Original z-stacks were cross-referenced to ensure that synaptic puncta correspond with MAP2+ dendrites. MAP2 somatodendritic area was determined from thresholded max-projected image. Synapses/MAP2 area was determined by dividing number of matched pre/postsynaptic puncta by MAP2 area for each imaged field. Percent of synaptic puncta apposed by post-synaptic partner was determined by dividing “Synapse Pre” objects by total “Pre No.” objects. N=3 or 4 experimental replicates, n≥30 imaging fields.

### Profile intensity analysis

Fiji’s Plot Profile tool was used to create plots of intensity values across distance for regions of interest for MAX-projected, multichannel image stacks. Intensity values were adjusted for background by subtracting the minimum intensity value and normalized to the maximum intensity value. The resulting intensity values were plotted vs. distance for each channel.

### Heterologous synapse intensity analysis

To determine Synapsin I/II or Synaptobrevin-2 intensity within Control or spastin KD axonal regions crossing NL1+ HEK cells, Fiji’s Thresholding tool was used to segment max-projection images and an 8-bit object mask based on the top 1% intensity was generated. Fiji’s Analyze Particles tool was used on the object mask redirected to the original image to determine intensity values. Individual object intensity values per HEK cell were averaged to generate HEK cell average, and cell averages were normalized to control replicate average. Normalized cell averages and experiment averages were plotted. N=3 experimental replicates, n=25 HEK cells encountering numerous crossing axons.

To determine spastin intensity within axonal regions crossing NL1+ or NL1-HEK cells, HEK cells were isolated and Fiji’s 3D Objects Counter tool was used to segment the image using Synaptobrevin-positive axonal regions. 3D measurements were redirected to the spastin channel. This volumetric analysis was performed to avoid interference of spastin intensity in adjacent HEK cells. Spastin intensity values within Synaptobrevin+ objects crossing HEK cells were averaged to generate HEK cell average, and cell averages were normalized to control replicate average. Normalized cell averages and experiment averages were plotted. N=3 experimental replicates, n=30 HEK cells encountering numerous crossing axons.

### SuperPlot data presentation and statistics

Experimental data were presented using the SuperPlot format [59], with color differentiating biological replicates. Replicate-matched data points (dots) and experimental averages (open circles) are displayed in the same color set to 50% and 100% opacity, respectively. Figure legends contain descriptions of the statistical test(s) used and specific *p*-values are displayed on the plots.

