## Supplemental Figures for "Spastin locally amplifies microtubule dynamics to pattern the axon for presynaptic cargo delivery"

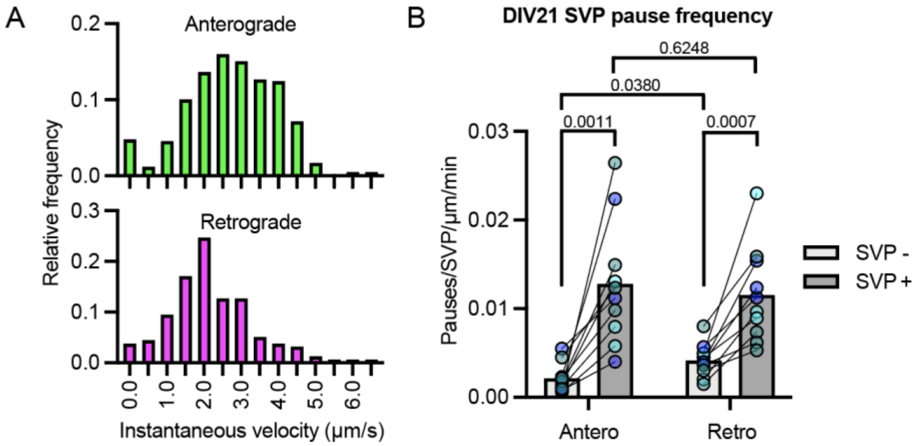


**Figure S1**

**(A)** Histograms of instantaneous velocities in μm/second for the anterograde (upper, magenta) and retrograde (lower, green) motility of SVPs in i$^3$Neurons at DIV35.

**(B)** Anterograde and retrograde SVP pause events in DIV21 i$^3$Neurons expressing mScarlet-Syp. Pauses are standardized for vesicle flux and distance in axonal areas lacking stable SVPs (SVP-, light gray) and populated by stable SVPs (SVP+, dark gray). Each of the paired data points represent the SVP- and SVP+ pause frequency within one axon. Reported *p*-values are from multiple paired t-test (between SVP+ and SVP- values) and from one-way repeated measures ANOVA (between anterograde and retrograde values).

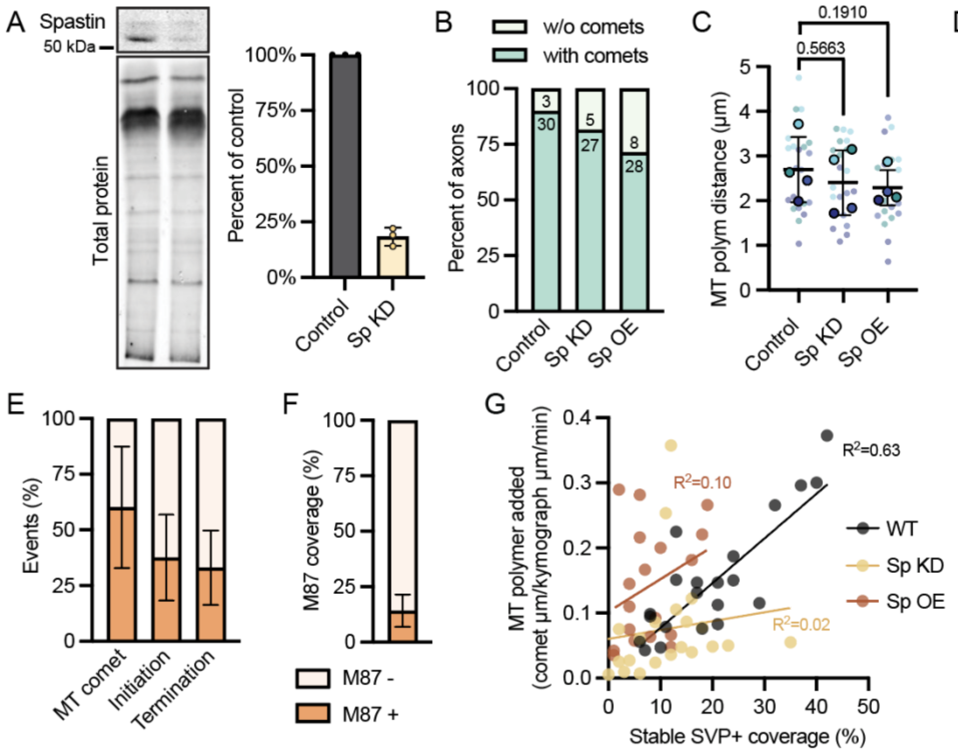


**Figure S2**

**(A)** Western blot of spastin level in CRISPRi non-targeting control (Control) and spastin CRISPRi-mediated knockdown (Sp KD) DIV21 $i^3$Neurons (upper panel). Data shown are normalized to total protein (lower panel); graph shows an average spastin level of 18±4% in CRISPRi spastin knockdown neurons relative to control neurons.

**(B)** Percent of imaged axons with clear MACF43 microtubule comets (with comets) and without clear comets (w/o comets) in Control, Sp KD, and Sp OE conditions. Number of axons in each category are provided on the plot.

**(C, D)** Microtubule polymerization distance (MT comet length; **C**) and duration (MT comet duration; **D**) for Control, Sp KD, and Sp OE. Errors bars represent the standard deviation of the replicate means and reported *p*-values are from one-way repeated measures ANOVA.

**(E)** Percent of microtubule comets, initiation events, and termination events that occur at M87-positive regions  (M87+ events/total events). Error bars represent the standard deviation of the data set.

**(F)** Axonal spastin SNAP-M87 coverage as percent of total axon distance (SNAP-M87 µm/total analyzed axon µm). Error bars represent the standard deviation of the data set.

**(G)** Stable SVP+ coverage as percent of total axon distance (stable SVP µm/total analyzed axon µm) versus new microtubule polymer added (comet µm/analyzed axon µm/min). Line of best fit and $R^2$ values are displayed for WT, Sp KD, and Sp OE data.

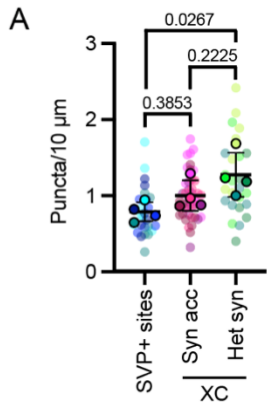


**Figure S3**

**(A)** Puncta density (puncta/10 µm of axon) of SVP+ sites in DIV21 axons (determined by live imaging), protosynaptic Syn accumulations (Syn acc; determined using XonaChip (XC) microfluidically isolated DIV14 axons), and heterologous synapses (Het syn; determined using XC-isolated DIV14 axons crossing NL1+ HEK cells).
